# Sequence-to-function deep learning decodes human *cis*-regulatory evolution

**DOI:** 10.64898/2026.08.24.746818

**Authors:** Riley J. Mangan, Nikitha Thoduguli, Dimitar Ivanov, Bessie Li, Krish Vasudev, Tom Zeerow, Jayashabari Shankar, Yuru Lin, Zunpeng Liu, Martin Wohlwend, Janet H.T. Song, Manolis Kellis

## Abstract

Deciphering the regulatory consequences of sequence divergence across human evolution is essential to understanding the molecular basis of human-specific traits and disease. Although millions of derived alleles distinguish humans from great apes, only a small fraction are likely to influence human-specific traits. Previous studies have focused on regions of elevated sequence divergence, assuming that rapid evolution reflects functional adaptation, yet individual high-impact regulatory mutations evade such scans. Here, we apply sequence-to-function deep learning to predict chromatin accessibility across modern human, archaic hominin, and great ape personalized genomes, identifying lineage-specific *cis*-regulatory elements (linCREs) across diverse cellular contexts. Compared to conserved elements, linCREs are shorter, less pleiotropic, less conserved, and enriched in neurodevelopmental pathways. Many linCREs occur in regions with limited sequence divergence that acceleration-based approaches would overlook. We validate lineage-specific enhancer activity through luciferase reporter assays and demonstrate that a single motif-generating derived allele nominated by model interpretability tools drives a hominin-specific neurodevelopmental enhancer.

## Main

Humans and other great apes are separated by vast phenotypic differences spanning cognition, bipedalism, metabolism, and life history^1,2^. As protein-coding mutations separating humans and great apes are sparse, it has been long suggested that regulatory mutations are central substrates of human-specific adaptations^3–5^. However, identifying the regulatory mutations underlying human-specific traits remains challenging, as millions of mutations have arisen on the human lineage, with the majority thought to have minimal functional or adaptive significance^6,7^.

To address this challenge, many studies have focused on divergence-based prioritization. This approach analyzes regions of rapid sequence divergence, measured as high concentrations of human-specific mutations, following the premise that anomalous rates of molecular evolution reflect functional divergence. Prominent examples identified using this approach include human accelerated regions^8^ (HARs) and human ancestor quickly evolved regions^9–11^ (HAQERs), which have been shown to function as human-specific enhancers, particularly in neurodevelopment^12,13^, and are enriched in neurodevelopmental and neuropsychiatric risk loci^9,14,15^.

While these divergence-based approaches have been highly valuable for identifying human-specific regulatory elements, we hypothesize that many phenotypically significant mutations do not occur in regions of rapid sequence divergence. Single mutations can be sufficient to alter the activity of regulatory elements that are otherwise unremarkable in their substitution rate, and such mutations are invisible to divergence-based scans^16,17^. However, comprehensive functional profiling across millions of derived variants in many distinct epigenomic contexts is infeasible, despite progress with massively parallel reporter assays (MPRAs) and CRISPR-based screens^18–21^.

Recent advances in sequence-to-function deep learning models, which are trained on vast epigenomic datasets to predict tissue-and cell-type-specific regulatory activity, enable the systematic evaluation of millions of mutations while circumventing direct experimental profiling^22,23^. While genomic deep learning models face challenges in predicting gene expression variation from personalized genomes^24–26^, they achieve reliable regulatory variant effect prediction when benchmarked against MPRAs^22^. Recent studies have begun applying these models to compare regulatory activity across species^27–31^.

Sequence-to-function models are uniquely valuable for studying the genomes of archaic hominins and ancient humans, where direct epigenomic profiling is impossible due to the absence of extant tissues. While ancient DNA studies have revealed historic selection^32,33^ and archaic introgression contributing to modern disease risk^34^, the regulatory consequences of variants identified by these approaches are largely uncharacterized. In this work, we apply sequence-to-function deep learning to predict chromatin accessibility across modern human, archaic hominin, and great ape personalized genomes, systematically identifying lineage-specific *cis*-regulatory elements (linCREs) across 100 epigenomic contexts. We show that linCREs recapitulate experimentally characterized human-ape regulatory and expression differences and independently recover previously defined divergence-based annotations. We find that linCRES are shorter, less pleiotropic, less conserved, and more tissue-specific than conserved elements, and are enriched in neurogenic and developmental pathways. Many linCREs exhibit regulatory divergence without elevated sequence divergence, escaping detection in acceleration-based scans. Finally, we leverage model interpretability to prioritize causal variants and their upstream regulators, validate lineage-specific enhancer activity by luciferase reporter assays, and demonstrate that a single motif-generating, human-derived variant is necessary to create a hominin-specific neurodevelopmental enhancer.

### Genome-wide prediction of lineage-specific *cis*-regulatory elements in human evolution

To identify candidate regulatory elements underlying lineage-specific phenotypes, we developed a strategy to systematically predict chromatin accessibility differences across hominins and great apes from sequence alone (**Fig. 1a**). We first generated diploid personalized genome assemblies for 17 individuals, including four high-coverage archaic hominins (one Denisovan, three Neanderthals), five modern humans, and eight great apes (4 chimpanzees, 1 bonobo, 2 gorillas, 1 orangutan), by reference-guided assembly of short-read sequencing libraries, producing 34 haplotypes aggregated into a 35-way multiple alignment (**Fig. 1a**, **Supplementary Notes 1-2**). We then used Enformer, a large pre-trained regulatory deep learning model^22^, to predict genome-wide chromatin accessibility from each haplotype across a curated panel of 100 epigenomic contexts selected to span the regulatory phenotypic space captured by the model (**Extended Data Fig. 1; Methods**), yielding 3,400 genome-wide accessibility prediction tracks.

**Figure 1:**
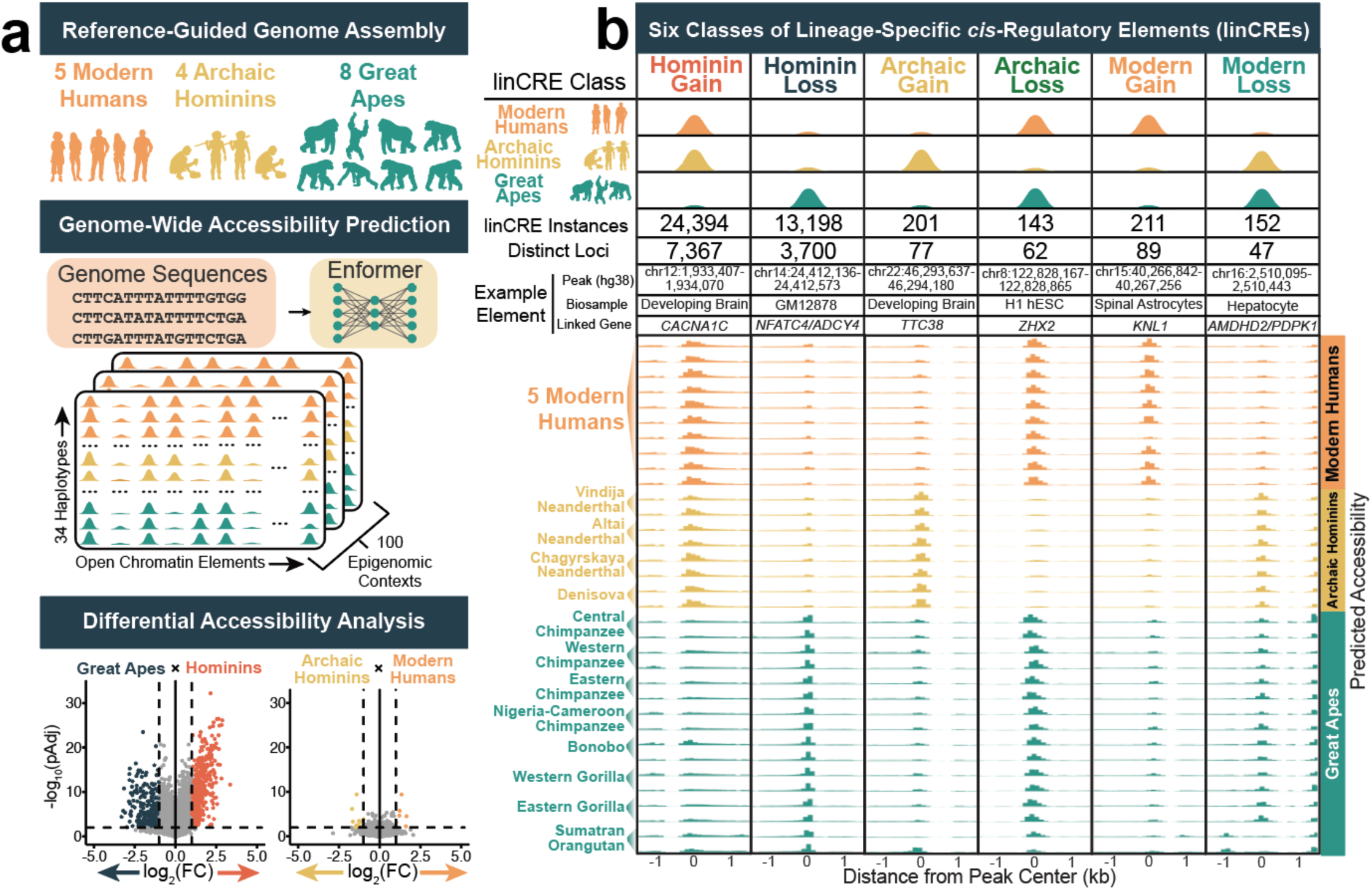
Sequence-based deep learning prediction of lineage-specific *cis*-regulatory elements in human evolution. (**a**) Identification of lineage-specific *cis*-regulatory elements (linCREs). We generated diploid personalized genomes across 17 hominin and great ape short read sequencing libraries through reference-guided assembly. We used Enformer^22^ to predict genome-wide chromatin accessibility profiles across 100 epigenomic contexts across all 34 haplotypes. We compared chromatin accessibility at open chromatin elements to identify linCREs. (**b**) Six classes of linCREs, defined by the lineage and direction of differential accessibility. Schematics at top illustrate accessibility profiles defining each class. Total linCRE instances and the number of distinct loci are reported for each class. Predicted accessibility across all 34 haplotypes in a 3kb window centered on a representative example element of each class are shown below, colored by lineage.

Because existing catalogs of regulatory elements derived from human biosamples may miss elements active selectively in archaic or great ape lineages, we identified open chromatin elements *de novo* from the prediction tracks themselves, identifying a high-confidence set of 161,981 distinct open chromatin elements (**Extended Data Fig. 2a-b; Methods; Supplementary Table 1**). These elements were strongly enriched for active enhancer and promoter chromatin states^35^, consistent with their interpretation as putative functional regulatory elements (**Extended Data Fig. 2c-f**).

We next tested each open chromatin element for differential predicted accessibility across haplotypes in two evolutionary comparisons: between hominins (including modern humans and archaic hominins) and great apes, and between modern humans and archaic hominins (**Fig. 1a; Extended Data Fig. 3**). We defined lineage-specific *cis*-regulatory elements (linCREs) as open chromatin elements with significantly differential predicted accessibility (FDR-adjusted P (pAdj) < 0.01, |log_2_ fold change| > 1) and named each class by the lineage and direction of change relative to the inferred ancestral state, using great apes as the outgroup (**Fig. 1b**).

In the hominin-great ape comparison, we identified 37,592 linCRE predictions across 11,050 distinct elements spanning diverse tissue and cell-type contexts (**Fig. 1b**; **Supplementary Tables 2-3**). These comprised 24,394 Hominin Gain predictions (7,367 distinct elements) and 13,198 Hominin Loss predictions (3,700 distinct elements), a ∼1.85-fold excess of gains over losses in predicted accessibility. This asymmetry is attributable to the structure of the comparison, as defining a Hominin Loss linCRE requires concordant accessibility across the included great ape species, which span a far greater evolutionary distance than the included hominins. This structure filters out chimpanzee-, bonobo-, gorilla-, or orangutan-specific elements, conservatively defining this set as hominin loss of ancestral regulatory activity.

Between modern humans and archaic hominins, we predicted 707 linCRE events across 235 distinct elements (**Fig. 1b**; **Supplementary Tables 4-7**). Polarizing each event against the great ape outgroup, we partitioned these into four single-lineage classes: Modern Gain (211 events in 89 elements), Modern Loss (152 events in 47 elements), Archaic Gain (201 events in 77 elements), and Archaic Loss (143 events in 62 elements). To assess specificity, we repeated the analysis after randomly permuting lineage labels across haplotypes, recovering no linCREs at our statistical threshold (**Extended Data Fig. 3a**).

To assess whether linCREs target loci with functional relevance, we linked each linCRE to candidate target genes using enhancer-gene linking predictions from the EpiMap resource^35^. Across linCRE classes, the majority of elements (52.1% to 64.5%) were linked to at least one putative target gene (**Extended Data Fig. 3b-c**, **Supplementary Table 8**). Notably, we observed a Hominin Gain-linCRE linked to *CACNA1C*, which encodes a calcium channel subunit, a major genetic risk factor for bipolar disorder, major depressive disorder, and schizophrenia^36^ (**Fig. 1b**). Human-specific structural variants within a *CACNA1C* intron have been previously shown to modulate regulatory activity, the neuronal response to stimulation^37^, and psychiatric disorder risk^38^, further implicating this locus in human-specific neurobiology. Other highlighted examples include an Archaic Gain-linCRE in the promoter of *TTC38*, an Archaic Loss-linCRE linked to the transcription factor *ZHX2*, and a Modern Gain-linCRE linked to *KNL1*, a kinetochore gene associated with microcephaly^39^ (**Fig. 1b**).

### Predicted linCREs recapitulate known human regulatory and expression changes

To assess the biological validity of our predictions, we tested linCREs for overlap with experimentally characterized sets of human-ape regulatory differences, sequence-based annotations of evolutionarily significant regions on the human lineage, and human-ape differentially expressed genes (**Fig. 2**). We first examined human-gained enhancers and promoters (HGE/Ps), elements identified by comparative ChIP-seq for H3K27ac and H3K4me2 across developing human, rhesus macaque, and mouse brains^40^. Hominin Gain-linCREs were strongly enriched for HGE/Ps (log_2_Enrich = 0.84, pAdj < 4.6 × 10^-37^), supporting these predictions as candidate regions for human-specific regulatory elements. Surprisingly, hominin Loss-linCREs were also modestly enriched for HGE/Ps (log_2_Enrich = 0.45, pAdj < 2.9 × 10^-5^), potentially reflecting repurposed regulatory elements with hominin gain of accessibility in one tissue and hominin loss in another. While HGE/Ps were ascertained in developing brains, overlap enrichments extended to linCRE sets identified in non-brain biosamples, suggesting that a subset of HGE/Ps and linCREs may exhibit regulatory divergence across broad epigenomic contexts (**Extended Data Fig. 4a, c**). We additionally examined human conserved deletions (hCONDELs), short human-specific deletions in otherwise conserved sequence whose regulatory consequences have been profiled by comparative MPRA^41^. With relatively few hCONDELs exhibiting species-specific regulatory activity, we were underpowered to detect set-level enrichments (**Fig. 2a**), though the directional pattern matched expectations, with Hominin Gain-linCREs trending toward enrichment at hCONDELs with human gain of MPRA activity and Hominin Loss-linCREs trending toward enrichment at hCONDELs with human-loss activity. This pattern persists when partitioning linCRE sets by biosample, though we note that these enrichments were driven by relatively few overlaps (**Extended Data Fig. 4b**).

**Figure 2:**
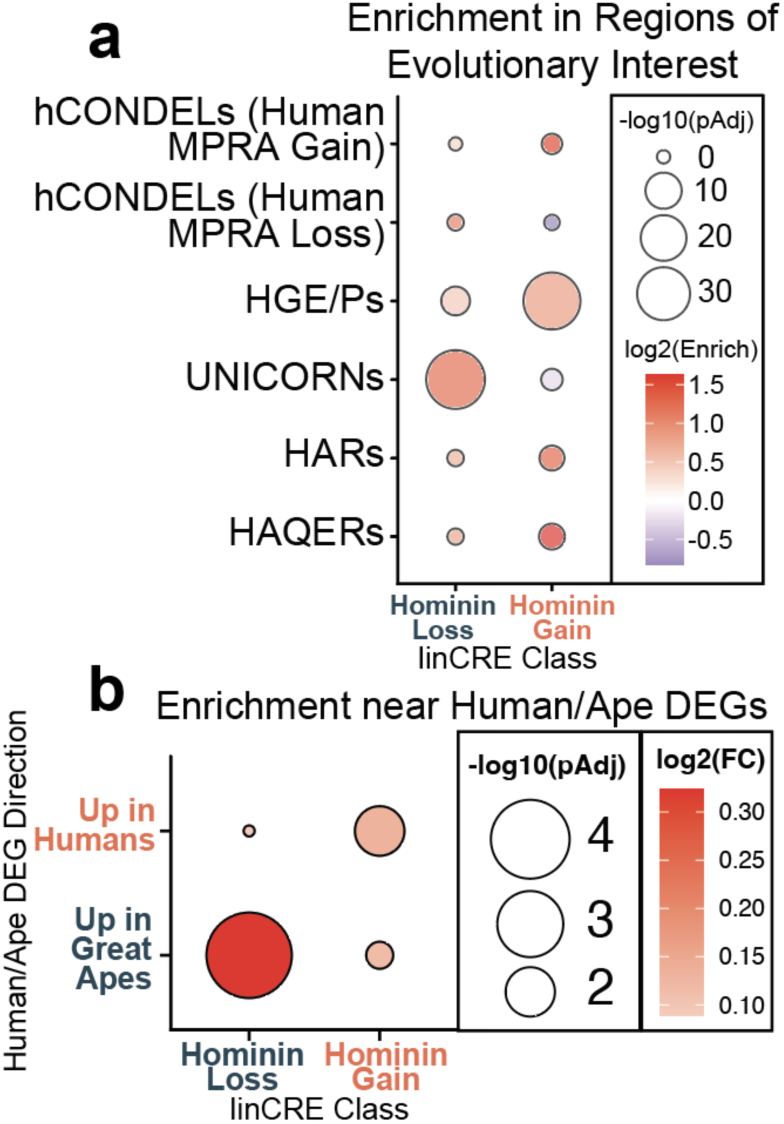
Predicted linCREs are enriched in evolutionary regions of interest and near human-ape differentially expressed genes. **(a)** Enrichment of predicted linCREs in previously defined regions of evolutionary interest. Hominin Gain elements were depleted in UNICORNs (unannotated intergenic constrained regions), whereas Hominin Loss elements were enriched, consistent with human-specific loss of ancestral regulatory function. Hominin Gain elements were enriched in both human accelerated regions (HARs) and human ancestor quickly evolved regions (HAQERs), while Hominin Loss elements were depleted in both. **(b)** Enrichment of linCREs near differentially expressed genes (DEGs) between humans and great apes in the middle temporal gyrus. Hominin Gain elements were enriched near genes upregulated in humans, while Hominin Loss elements were enriched near genes upregulated in apes.

Turning to sequence-based annotations of human evolutionary significance, we first examined unannotated intergenic constrained regions (UNICORNs), regions lacking regulatory annotations in humans despite deep mammalian conservation^42^. Notably, UNICORNs were nominally depleted from Hominin Gain-linCREs (log_2_Enrich = -0.25, pAdj = 0.054) and significantly enriched in Hominin Loss-linCREs (log_2_Enrich = 1.2, pAdj < 4.4 × 10^-40^). This finding suggests that sequence conservation in UNICORNs may reflect ancestral regulatory activity lost in the human lineage, consistent with their lack of functional annotation in human datasets. Predicted Hominin Gain-linCREs were also significantly enriched in HARs^14^ (log_2_Enrich = 1.3, pAdj < 0.0028) and HAQERs^9^ (log_2_Enrich = 1.63, pAdj < 0.001), demonstrating that our predictions independently recover previous divergence-based annotations of human regulatory innovation. Finally, we asked whether predicted regulatory divergence is reflected in lineage-specific gene expression. We tested for overlap between linCREs and genes differentially expressed between humans and great apes (chimpanzees and gorillas) in the middle temporal gyrus^43^, treating each gene as the genomic interval extending ± 50kb from the transcription start site. The union set of Hominin Gain-linCREs were significantly enriched near genes upregulated in humans (log_2_Enrich = 0.13, pAdj < 9.8 × 10^-3^), and the union set of Hominin Loss-linCREs were enriched near genes upregulated in chimpanzees and gorillas (log_2_Enrich = 0.32, pAdj < 2.6 × 10^-5^) (**Fig. 2b**). This directional concordance between predicted regulatory divergence and lineage-specific transcriptional programs supports a contribution of predicted linCREs to human-ape gene expression divergence.

### Predicted linCREs bear signatures of evolutionarily malleable regulatory elements

To understand what distinguishes regulatory elements that diverge between lineages from those with conserved accessibility, we characterized regulatory states, constraint, and functional associations of linCREs. Both Hominin Gain and Hominin Loss-linCREs were strongly depleted for promoter (TssA) chromatin states relative to elements with conserved accessibility (**Fig. 3a**, 30.7% and 19.7% vs. 41.9%; p < 10^-82^ and p < 10^-176^, respectively), identifying linCREs as predominantly distal enhancers. As enhancers evolve more freely than the promoters they regulate^44^, this depletion is consistent with enhancers as the principal substrate of human regulatory divergence. Hominin Loss-linCREs were further enriched for elements falling outside any annotated regulatory state, relative to both Hominin Gain-linCREs and conserved elements (**Fig. 3b**, p < 4.3 × 10^-6^ and p < 7.0 × 10^-13^, respectively), as expected if these elements reflect ancestral regulatory activity lost in humans and therefore absent from human-derived annotations, consistent with the enrichment of Hominin Loss-linCREs in UNICORNs (**Fig. 2a**).

**Figure 3:**
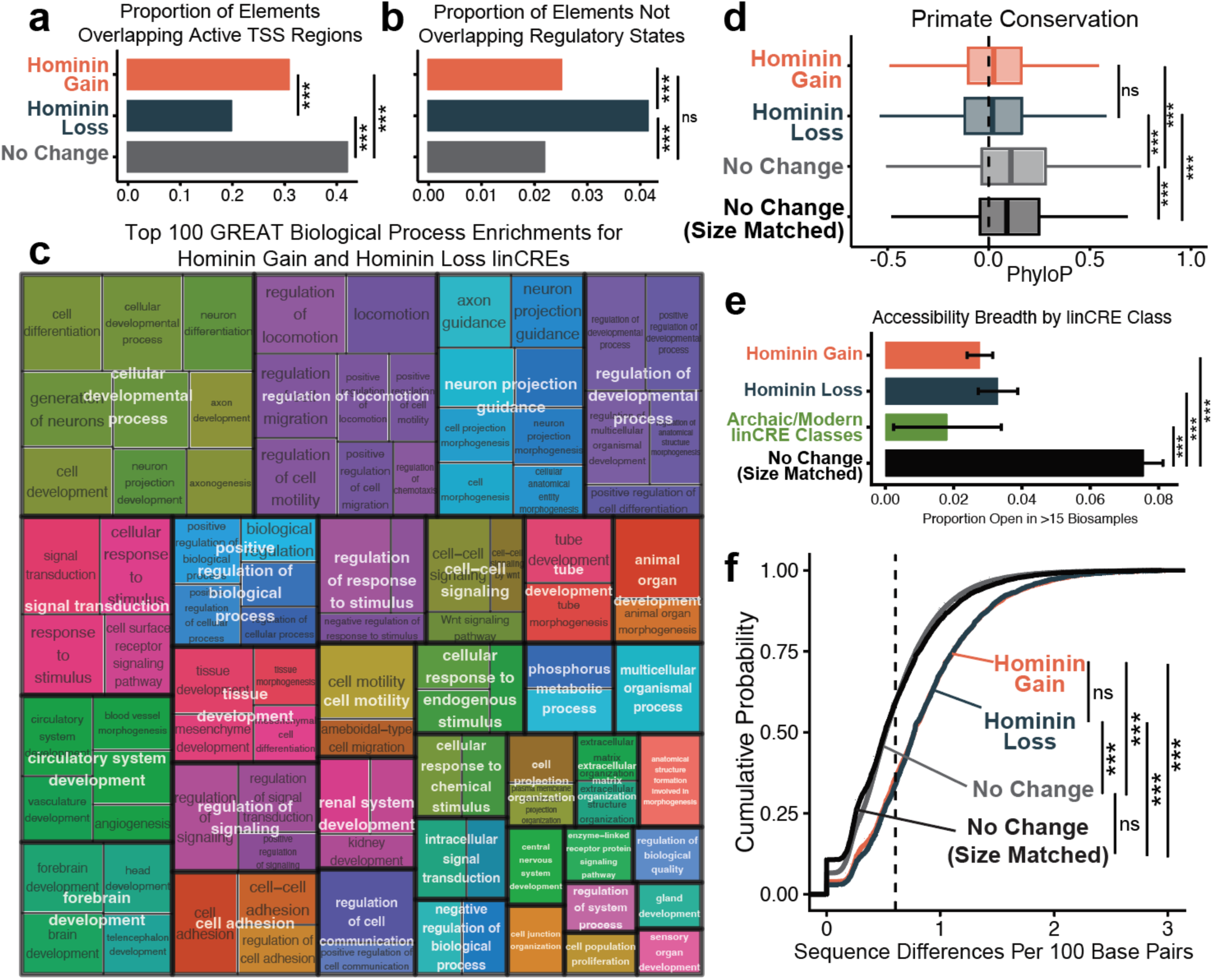
linCREs are evolutionarily malleable elements that diverge with limited sequence change. (**a**) Proportion of open chromatin elements overlapping active transcription start site (TssA) chromatin states. Predicted linCREs are significantly depleted for promoter annotations relative to elements with conserved accessibility (Fisher’s exact). (**b**) Proportion of elements not overlapping any regulatory ChromHMM states. Hominin Loss-linCREs are significantly more likely to fall outside annotated regulatory states, consistent with human-specific loss of ancestral regulatory function. (**c**) Semantic clustering of gene ontology enrichments for Hominin Gain and Hominin Loss-linCREs. Top enriched biological process terms included neurogenic and developmental pathways. (**d**) Primate conservation scores across open chromatin elements. Hominin Gain and Hominin Loss-linCREs exhibit less sequence conservation than elements with conserved accessibility. (**e**) Accessibility breadth by linCRE class, quantified as the proportion of elements open in >15 biosamples. linCREs exhibit significantly less breadth when compared to the set of CREs with conserved accessibility (Fisher’s exact). (**f**) Local sequence divergence (sequence differences per 100 bp) across element classes. The dotted line indicates the genome-wide average divergence rate in a 100 bp window. (*** p < 0.001, ns = not significant)

Both Hominin Gain and Hominin Loss-linCREs were most prominently enriched near genes governing development and neurogenesis (**Fig. 3c**; **Supplementary Table 9**). Notably, Hominin Gain and Hominin Loss-linCREs converge on many of the same developmental and neurogenic pathways, suggesting that these gene programs are recurrent targets of *cis*-regulatory turnover in human evolution (**Extended Data Fig. 5a-c, Supplemental Tables 10-11**). linCREs distinguishing modern humans from archaic hominins are most enriched near pathways related to neurogenesis, response to hypoxia, and chemotaxis (**Extended Data Fig. 5d, Supplemental Table 12**).

We next asked whether linCREs show the evolutionary properties expected of evolutionarily malleable regulatory elements, finding four convergent signals. First, linCREs are shorter in length than conserved elements, even when accounting for length differences between promoters and distal enhancers (**Extended Data Fig. 6a-d**). Second, linCREs were less constrained than conserved elements, with lower phyloP scores across both primate and mammalian alignments, a difference that persisted after size-matching and after stratifying by chromatin state (**Fig. 3d; Extended Data Fig. 6e-h**). Third, linCREs were active in fewer biosamples, indicating more restricted regulatory function (Measured as the proportion of elements open in > 15 Enformer tracks. Hominin Gain-linCRE: 2.8%, Hominin Loss-linCRE: 3.3%, Combined Modern/Archaic linCRE classes: 3.4%, Size-matched Conserved CREs: 8.1%, **Fig. 3e**). Fourth, fewer variants within linCREs exhibited significant horizontal pleiotropy than in conserved elements, indicating more restricted impact on human disease phenotypes^45^ (**Extended Data Fig. 6i,j**). As mutations in broadly active, strongly conserved, and highly pleiotropic elements are most likely to be deleterious, these features indicate that regulatory divergence occurs in the most evolutionarily malleable portions of the genome, offering orthogonal support that our deep learning predictions capture true regulatory turnover rather than technical noise.

### Functional divergence can occur without elevated sequence divergence

A central premise of our approach is that phenotypically consequential regulatory changes need not coincide with rapid sequence divergence. To test this directly, we counted human-chimpanzee-ancestor sequence differences^9^ (SNPs and short INDELs) per 100 base pairs within each element class. As expected, linCREs carried more sequence divergence than conserved elements (p < 2 × 10^-16^, pairwise t-test; **Fig. 3f**). Notably, this effect was modest in magnitude: 34.9% of Hominin Gain and Hominin Loss-linCREs fell below the genome-wide average human-ape divergence. Thus, although functional divergence is associated with elevated sequence change on average, a substantial fraction of lineage-specific regulatory elements would be invisible to divergence-based scans, supporting our premise that functional innovations can arise from a single or small number of high-impact mutations in regulatory elements.

### Identifying causal variants underlying lineage-specific accessibility

We next leveraged the interpretability of sequence-to-function models to prioritize causal variants and infer the upstream transcription factor regulators underlying lineage-specific regulatory function. We applied *in silico* saturation mutagenesis (ISM), in which every possible single nucleotide variant is introduced into a sequence to measure its predicted effect on accessibility. We first performed ISM experiments with Enformer to nominate candidate causal variants, then re-evaluated each prioritized variant with AlphaGenome, a recent successor model to Enformer^23^. Across all tested loci, the two models yielded concordant predictions for underlying regulatory mechanisms.

For many linCREs, ISM prioritized a single variant as sufficient to explain lineage-specific accessibility, often corresponding to lineage-specific gains and losses of a transcription factor binding site (TFBS) identified as important to the accessibility prediction. First, at a brain Hominin Gain-linCRE linked to *GRIN3B*, the derived allele creates a novel E2F1/2 motif (**Extended Data Fig. 7a-b**). Notably, this substitution simultaneously acts as a missense mutation within an exon of the overlapping gene *ATP8B3*, with the ancestral allele retained at very low frequency in modern humans (MAF = 8.43 × 10^-5^). Second, at a Modern Gain-linCRE active in astrocytes linked to *KNL1*, the derived allele creates a FOS motif, consistent with recent findings that variants in JUN/FOS binding sites frequently drive species-specific enhancer activity in human neurons^46^ (**Extended Data Fig. 7c-d**). This site remains polymorphic in modern humans (rs12439947, MAF ∼0.17), and a single human haplotype carrying the ancestral allele showed markedly reduced predicted accessibility (**Extended Data Fig. 7c**). While *KNL1* is essential for neural progenitor proliferation, and disruption causes microcephaly^39^, phenotypic consequences of its altered regulation in astrocytes are unclear. Notably, *KNL1* carries two missense mutations separating modern humans from most Neanderthals, suggesting this locus may have been subject to natural selection in recent human evolution^47^. Third, at a Hominin Gain-linCRE in an intron of *KCNS1*, a single derived allele (nearly fixed, ancestral allele frequency = 3.8 × 10^-6^) creates a hominin-specific CTCF motif that both Enformer and AlphaGenome identify as the driver of accessibility and CTCF binding (**Extended Data Fig. 7e-g**).

Beyond single-lineage changes, we identified 17 loci where an element predicted as a Hominin Gain-linCRE in one biosample overlapped an element predicted as a Hominin Loss-linCRE in another, a pattern suggestive of tissue-specific repurposing of ancestral regulatory elements (**Extended Data Fig. 8a**). One such element in the 3’ UTR of *LAD1* is linked to cardiac troponin genes *TNNT2* and *TNNI1* and harbors a Hominin Gain-linCRE active in the developing heart that partially overlaps a Hominin Loss-linCRE active in hepatocytes (**Extended Data Fig. 8b**). ISM prioritized two distinct causal variants: a derived allele creating a human-specific Nuclear Factor I motif underlying the heart-specific gain, and a separate derived allele ablating an ancestral JUNB/FOSB motif underlying the hepatocyte-specific loss (**Extended Data Fig. 8c-f**). Both derived alleles appear fixed in modern humans, as neither ancestral allele is observed in dbSNP.

### Reporter assays validate predicted lineage-specific regulatory activity

We next sought to validate these *in silico* predictions through luciferase reporter assays in neural stem/progenitor cells (NSPCs). We selected candidates from 55 Hominin Gain and 23 Hominin Loss-linCREs predicted in the Enformer NSPC context, prioritizing elements identified as linCREs across multiple brain-related biosamples and linked to annotated target genes, particularly those that are differentially expressed between human and great ape brains^43^ (**Supplemental Table 13**, **Methods**). We amplified orthologous sequences from human and chimpanzee genomic DNA, successfully cloning 12 human/chimpanzee ortholog pairs. We additionally included three Hominin Gain elements predicted in the Enformer developing brain context linked to neurodevelopmental genes of interest (*CACNA1C*, *GRID1*, and *GRIN3B*) yielding 15 ortholog pairs in total (**Supplemental Table 13**). We cloned each ortholog upstream of luciferase and measured reporter activity across four NSPC lines derived from induced pluripotent stem cells (iPSCs), whose differentiation we validated through marker gene expression (**Extended Data Fig. 9a**). We derived NSPCs from two human and two chimpanzee iPSC lines, enabling us to disentangle sequence-encoded *cis*-regulatory differences from differences attributable to species-specific *trans*-regulatory environments (**Fig. 4a, Supplementary Table 14**).

**Figure 4:**
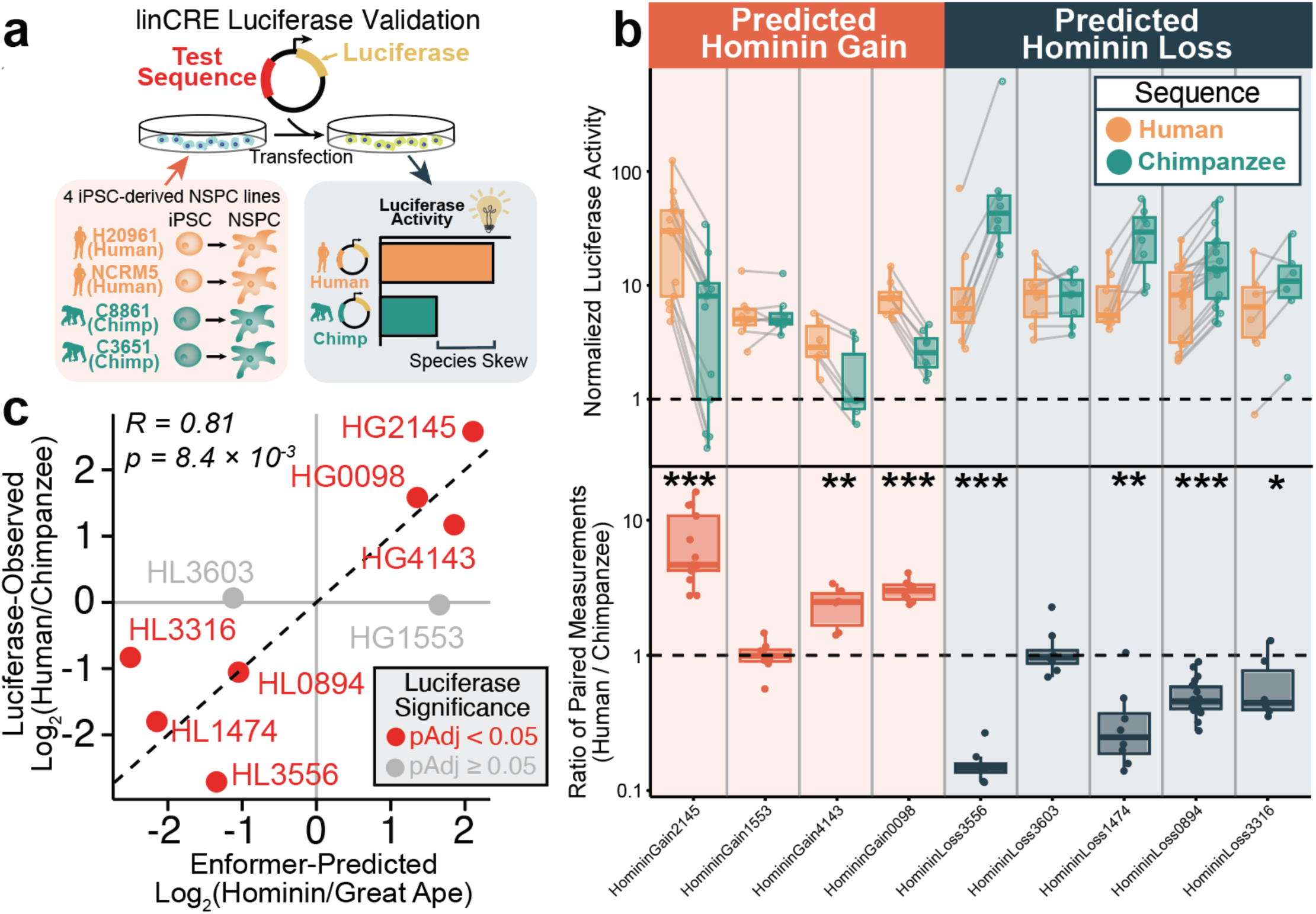
Functional validation of predicted linCREs by luciferase reporter assay in neural stem/progenitor cells. (a) Experimental design for linCRE luciferase validation. Human and chimpanzee orthologous sequences for predicted linCREs were cloned upstream of luciferase and transfected into iPSC-derived neural stem/progenitor cells (NSPCs) from four lines (two human: H20961, NCRM5; two chimpanzee: C8861, C3651). We compared luciferase activity between orthologs to quantify species-specific regulatory activity. (b) Luciferase measurements for nine predicted linCREs grouped by predicted class (left Predicted Hominin Gain, right Predicted Hominin Loss). Top, normalized luciferase activity for human (orange) and chimpanzee (teal) sequences, with paired measurements linked by gray lines. Bottom, ratio of paired human/chimpanzee measurements. The dashed line indicates no human/chimpanzee differences; observations above the dashed line indicate greater luciferase activity from the human sequence, observations below the dashed line indicate greater luciferase activity from the chimpanzee sequence. Asterisks denote elements with significant human-chimpanzee differences (One-sided t-test on log_2_ ratio, BH-adjusted. *: p < 0.05, **: p < 0.01, ***: p <0.001). (c) Concordance between Enformer-predicted log_2_(hominin/great ape) accessibility and luciferase-observed log_2_(human/chimpanzee) activity across nine active elements (Pearson R = 0.81, P = 8.4 × 10^-3^). Points are colored by luciferase significance (red: pAdj < 0.05; gray: not significant).

We first asked whether each element functioned as an enhancer in cultured NSPCs, testing whether the human or chimpanzee ortholog drove reporter activity above background. 9 of the 15 tested elements showed significant enhancer activity in at least one species (**Extended Data Fig. 9b**). None of the three developing brain elements were active, consistent with the fact that these elements were not predicted as linCREs in NSPCs. Among elements prioritized specifically in the Enformer neural stem/progenitor cell context, 9 of 12 were active. Because reporter measurements spanned multiple cell lines and plate batches, we quantified species-specific activity using paired measurements between human and chimpanzee orthologs within each plate, controlling for batch effects. Of the nine active elements, seven exhibited significant differences in activity between the human and chimpanzee sequences. For all seven of these elements, the observed direction of lineage-specific enhancer activity matched the model prediction: three Hominin Gain-linCREs (HomininGain2145, HomininGain4143, and HomininGain0098) showed significantly greater activity from the human sequence, and four Hominin Loss-linCREs (HomininLoss3556, HomininLoss1474, HomininLoss0894, and HomininLoss3316) showed significantly greater activity from the chimpanzee sequence (**Fig. 4b**). Enformer-predicted accessibility fold changes were strongly correlated with observed luciferase activity ratios (Pearson R=0.81, p = 8.4 × 10^-3^, **Fig. 4c**). Luciferase activity was positively correlated across all pairs of cell lines, with modestly higher concordance within species than across species (**Extended Data Fig. 9e-f**). Nonetheless, species differences in reporter activity were largely preserved when analyses were stratified to observations in human or chimpanzee NSPC lines, indicating that *trans* differences do not confound our evaluation of lineage-specific enhancer activity (**Extended Data Fig. 9c-f**). Together, these results demonstrate through an orthogonal functional assay that sequence-based deep learning effectively prioritizes lineage-specific *cis*-regulatory divergence.

### A motif-generating derived allele creates a hominin-specific neurodevelopmental enhancer

We next dissected a single high-confidence locus to resolve the causal basis of its lineage-specific regulatory activity at base-pair resolution. HomininGain2145, an element exhibiting hominin-specific activity in neural stem/progenitor cells (**Fig. 4b-c**, **Fig. 5a**), is linked to *GNB5*^35^, a gene in which biallelic loss-of-function mutations cause an autosomal recessive syndrome whose neurological features include intellectual disability, language deficits, and developmental delay^48^. *GNB5* was also identified as differentially expressed between humans and great ape brains in a comparative single-cell study^43^, consistent with a lineage-specific regulatory change at this locus. ISM prioritized a single derived variant (labeled variant 1) as the primary determinant of predicted accessibility divergence, with reciprocal predicted variant effects relative to the human (hg38) and chimpanzee (panTro6) reference genomes (**Fig. 5b**), a prediction supported by AlphaGenome (**Extended Data Fig. 10a**). Notably, the derived allele at variant 1 creates a human-specific serum response factor (SRF) TFBS, providing a candidate molecular mechanism for the hominin-specific gain in accessibility (**Fig. 5b**). While the ancestral C allele is absent from dbSNP, indicating fixation, a rare secondary polymorphism is present at this position in modern humans (rs142420474, T → A; MAF ∼ 0.003). However, the A allele remains compatible with the SRF motif and is predicted by Enformer to largely maintain the hominin-gained accessibility (**Fig. 5b**). We next generated mutant luciferase constructs to test whether this single variant is causal for the observed regulatory divergence (**Fig. 5c**). Reverting the derived allele in the human sequence (Hu_T→C_) significantly reduces luciferase activity relative to the wildtype human sequence (Hu) (Hu / Hu_T→C_, pAdj = 8.63 × 10^-5^, **Fig. 5d-e**), effectively returning it to chimpanzee levels (Hu_T→C_ / Ch; pAdj = 0.261). Conversely, introducing the derived allele into the chimpanzee sequence (Ch_C→T_) significantly increased luciferase relative to the wildtype chimpanzee sequence (Ch) (Ch_C→T_ / Ch, pAdj = 0.014, **Fig. 5d-e**). However, reporter activity from the chimpanzee mutant construct remained significantly lower than the wildtype human sequence (Hu / Ch_C→T_; pAdj = 8.63 × 10^-5^). Together, these observations identify a single derived allele as necessary and sufficient to confer hominin-specific gain of enhancer activity at this locus.

**Figure 5:**
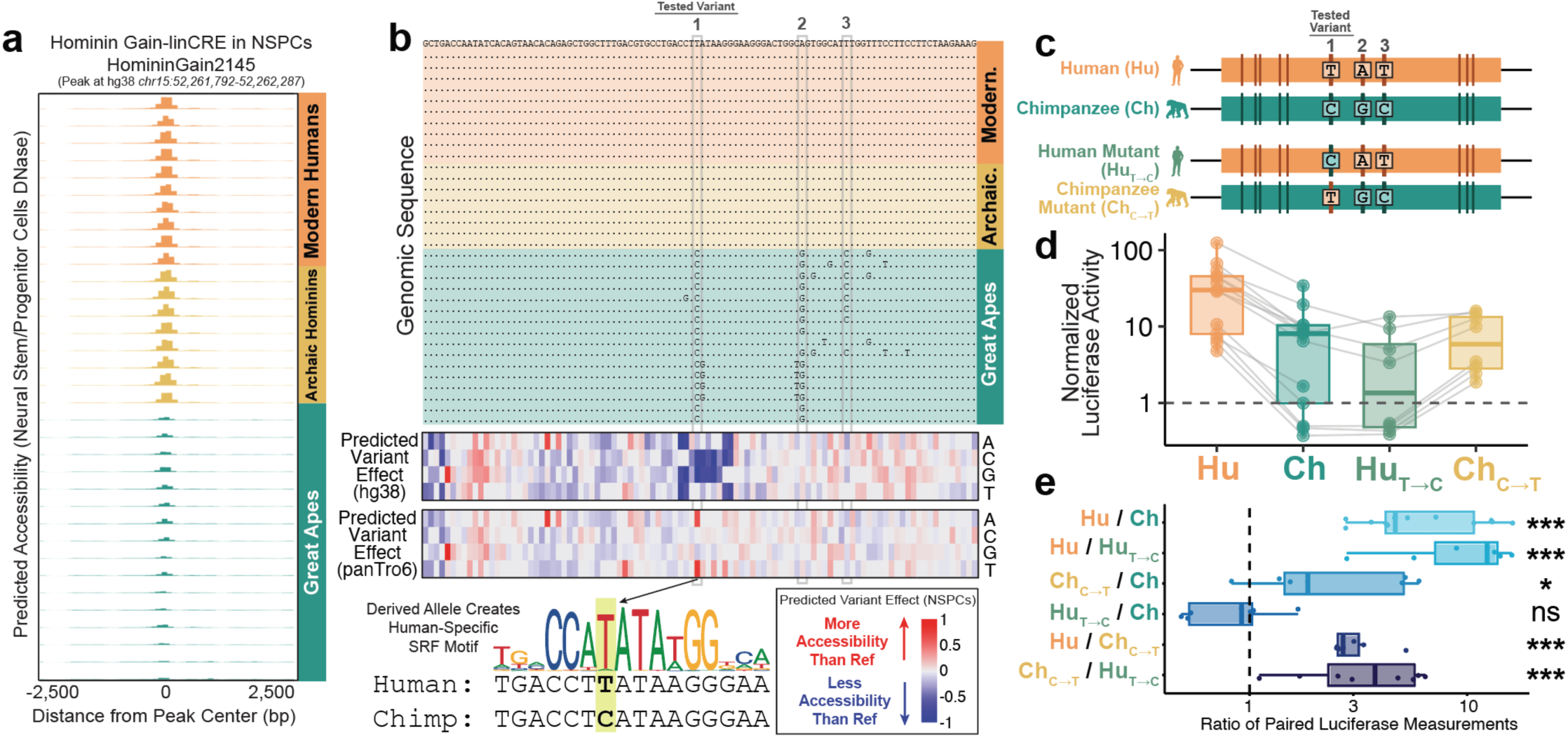
A motif-generating derived allele drives a hominin-specific neurodevelopmental enhancer. (a) Predicted chromatin accessibility across all 34 haplotypes in neural stem/progenitor cells (NSPC DNase) at HomininGain2145, showing hominin-specific predicted accessibility, colored by lineage. (b) Top: genomic sequence across 34 haplotypes in a 100bp segment of HomininGain2145 containing three segregating sites (Variant 1-3). Middle: in silico saturation mutagenesis (ISM) matrices computed relative to both the human (hg38) and chimpanzee (panTro6) reference genomes, which reciprocally identify variant 1 as the primary determinant of predicted accessibility. Bottom: the derived allele at variant 1 (T, versus ancestral C) creates a human-specific SRF transcription factor binding site. (c) Luciferase construct design. Human (Hu) and chimpanzee (Ch) reference sequences and reciprocal single-variant mutants at variant 1, including Hu_T→C_, the human sequence carrying the chimpanzee allele at variant 1, and Ch_C→T_, the chimpanzee sequence carrying the human allele at variant 1. (d) Normalized luciferase activity for the four constructs in NSPCs. Paired measurements are linked by gray lines. (e) Ratio of paired luciferase measurements for construct comparisons. Variant 1 is necessary and sufficient for hominin-gained enhancer activity (t-test on log_2_ ratio, BH-adjusted. *: p < 0.05, **: p < 0.01, ***: p < 0.001, ns = not significant).

## Discussion

We present a deep learning atlas of *cis*-regulatory evolution, predicting lineage-specific regulatory elements from personalized genome sequences from modern humans, archaic hominins, and great apes. By comparing chromatin accessibility tracks predicted for orthologous sequences across 100 epigenomic contexts, we defined lineage-specific *cis*-regulatory elements (linCREs) spanning both hominin-great ape divergence and the more recent split between modern and archaic hominins. These predictions are enriched in experimentally characterized human-ape regulatory innovations and near human-ape differentially expressed genes, and are enriched in divergence and conservation-based evolutionary annotations such as HARs^8^, HAQERs^9^, and UNICORNs^42^.

For two decades, the discovery of evolutionarily significant noncoding regions has relied primarily on identifying local elevations in sequence divergence rates. Here, we shift the analytical paradigm from counting mutations to prioritizing derived alleles with outsized regulatory impact, leveraging deep learning models trained to interpret cell-type-specific regulatory grammars to provide an orthogonal approach for discovering functional evolutionary change. While we observe significant co-localization between linCREs and HARs/HAQERs, we find that most linCREs exhibit only modest sequence divergence, suggesting that major regulatory innovations may be mediated by single high-impact mutations that remain invisible to traditional divergence-based scans^49^. Through reporter assays, we confirm the high efficacy of this approach to predict lineage-specific regulatory elements. Furthermore, we illustrate how deep learning interpretability methods can be leveraged to dissect the precise molecular mechanisms underlying human-specific enhancer function. By swapping a human-derived and ancestral variant at HomininGain2145, we demonstrate that a single model-prioritized variant creating an SRF binding site confers a hominin-specific neurodevelopmental enhancer. Both Hominin Gain-and Hominin Loss-linCREs share strong enrichments near neurogenic and developmental genes, pointing to these programs as recurring targets of *cis*-regulatory innovation in human evolution. Several linCREs implicate genes with established roles in human neurobiology and disease, including *GNB5*, *CACNA1C*, and *KNL1*, suggesting a complex interplay between ancient regulatory evolution and modern disease susceptibility^15^.

Recent concurrent studies have similarly leveraged sequence-to-function models to predict human-gained regulatory elements using human and great ape reference genomes, either charting evolutionary histories in the developing cerebellum across mammals^30^ or identifying regions of increased accessibility across a human single-cell atlas^29^. While prediction sets are enriched for each other across studies, most individual predictions are study-specific, reflecting differences in training data, deep learning architectures, and sampled epigenomic contexts (**Supplementary Note 3**).

Beyond extant species, sequence-to-function prediction provides opportunities to address limitations in archaic variant interpretation. In the absence of extant tissues, archaic regulatory biology is largely inaccessible to experimental profiling. Notable exceptions include ancient DNA methylation, which is largely restricted to skeletal epigenomic contexts^50^ and analysis of introgressed haplotypes, which is restricted to archaic variation retained in modern populations^34^. Recent work applying sequence models to predict divergence in 3D chromatin organization between modern and archaic hominins illustrate the broader promise of this approach^51^, as does our current work, nominating hundreds of elements with predicted modern-archaic functional divergence.

The sequence and evolutionary features distinguishing linCREs from conserved elements align with the conceptual model where young regulatory elements emerge from neutral sequence as short, weakly conserved ‘proto-enhancers’ that accumulate transcription factor binding sites, growing in length, constraint and pleiotropy over time^49,52,53^. Beyond *de novo* gains and losses, we identified several elements repurposed across cellular contacts, predicted as hominin gains in one biosample and losses in another. At one such element linked to the cardiac troponin genes *TNNT2* and *TNNI1*, distinct derived variants independently drive heart-specific gain and hepatocyte-specific loss of predicted activity. This predicted regulatory repurposing towards cardiac regulation is notable in the context of comparative anatomical studies demonstrating that the human left ventricle has evolved derived features supporting endurance activity characteristic of hunter-gatherer subsistence strategies, in contrast to the short bursts of resistance-based activity characteristic of great apes^54^. However, these repurposed elements are relatively rare in our prediction set, indicating that linCREs predominantly reflect *de novo* gains or losses of activity. More broadly, the relative contributions of gain, loss, and repurposing to regulatory evolution remains unclear^55^.

Our study has several limitations that invite future work. First, our reliance on short-read sequencing libraries excludes repetitive sequences that may harbor evolutionarily significant regulatory elements. While telomere-to-telomere assemblies address this limitation for extant humans and great apes^10,11^, they are infeasible for naturally fragmented ancient DNA libraries. We anticipate that pangenome-based reconstruction of introgressed archaic haplotypes may offer a path forward^56^. Second, while our sequence-based approach has the advantage of isolating *cis*-regulatory effects over epigenomic comparisons that conflate *cis* and *trans* influences, we do not model *trans*-regulatory divergence, whose contribution to primate evolution is increasingly appreciated^57,58^. Future models trained directly on great ape epigenomes or human-chimpanzee tetraploid systems may address this. Third, linking regulatory divergence to gene expression and organismal phenotypic transitions remains an open challenge given the known difficulty of predicting expression-altering variants directly from personalized genomes^24–26^. Furthermore, recent studies highlighting the disconnect between molecular regulatory variation and complex trait heritability suggest that predicted linCREs likely represent a heterogeneous collection of both selectively neutral regulatory turnover events and functionally consequential drivers of human adaptation^59^. Future experimental work to evaluate linCREs through CRISPRi perturbations^12^, massively parallel reporter assays^60^, and transgenics^13,61^ will allow us to connect linCREs to their molecular and cellular consequences, ultimately supporting a mechanistic understanding of the sequence changes that underlie human-specific traits and disease.

## Supplemental Tables

Supplemental Table 1. Genomic locations of 161,981 open chromatin elements analyzed in this study, reported on hg38 coordinates.

Supplemental Table 2. Genomic coordinates (hg38) of 7,367 predicted Hominin Gain-linCREs.

Supplemental Table 3. Genomic coordinates (hg38) of 3,700 predicted Hominin Loss-linCREs.

Supplemental Table 4. Genomic coordinates (hg38) of 77 predicted Archaic Gain-linCREs.

Supplemental Table 5. Genomic coordinates (hg38) of 62 predicted Archaic Loss-linCREs.

Supplemental Table 6. Genomic coordinates (hg38) of 89 predicted Modern Gain-linCREs.

Supplemental Table 7. Genomic coordinates (hg38) of 47 predicted Modern Loss-linCREs.

Supplemental Table 8. Predicted target genes for each linCRE across 31 tissues from the EpiMap enhancer-gene linking resource.

Supplemental Table 9. GREAT ontology analysis for the combined set of Hominin Gain and Hominin Loss-linCREs.

Supplemental Table 10. GREAT ontology analysis for Hominin Gain-linCREs.

Supplemental Table 11. GREAT ontology analysis for Hominin Loss-linCREs.

Supplemental Table 12. GREAT ontology analysis for the combined set of Archaic Gain, Archaic Loss, Modern Gain, and Modern Loss linCREs.

Supplemental Table 13. Details on 15 linCREs tested in luciferase assays.

Supplemental Table 14. Raw experimental results from luciferase assays.

Supplemental Table 15. Oligonucleotides used for luciferase cloning and mutagenesis.

## Acknowledgements

Research reported in this publication was supported by the National Institute of General Medical Sciences of the National Institutes of Health under Award Number “T32 GM007748” (R.J.M.) and by the National Institute of Mental Health of the National Institutes of Health under Award Number R00MH136290 (J.H.T.S). We would like to acknowledge The Bauer Core Facility at Harvard University for providing us access to the BioTek Synergy Neo2 plate reader used for the luciferase assays. We would also like to acknowledge members of the Song and Capellini Labs at the Department of Human Evolutionary Biology at Harvard University, members of the Kellis lab at MIT CSAIL, as well as Craig B. Lowe and members of his lab in the Department of Molecular Genetics and Microbiology at Duke University for their thoughtful comments and feedback.

## Author Contributions

RJM, JHTS, and MK designed the study. RJM, NT, BL, KV, JS, YL, ZL, and MW performed computational analyses. DI and TZ performed and analyzed luciferase reporter assays. JHTS and MK supervised and funded research. All authors reviewed, edited, and approved the final manuscript.

## Data and Code Availability

Gonomics programs described in this manuscript, including the ANCoRA personalized genome assembly and simulation-based validation framework, are available from the Gonomics repository and accessible at http://github.com/vertgenlab/gonomics.

Code used for analysis and to visualize data is available at https://github.com/rimangan/evolutionDeepLearning. These materials, as well as raw accessibility prediction matrices and genome alignments, are available at Zenodo at https://doi.org/10.5281/zenodo.22047727.

## Extended Data Figures

**Extended Data Figure 1:**
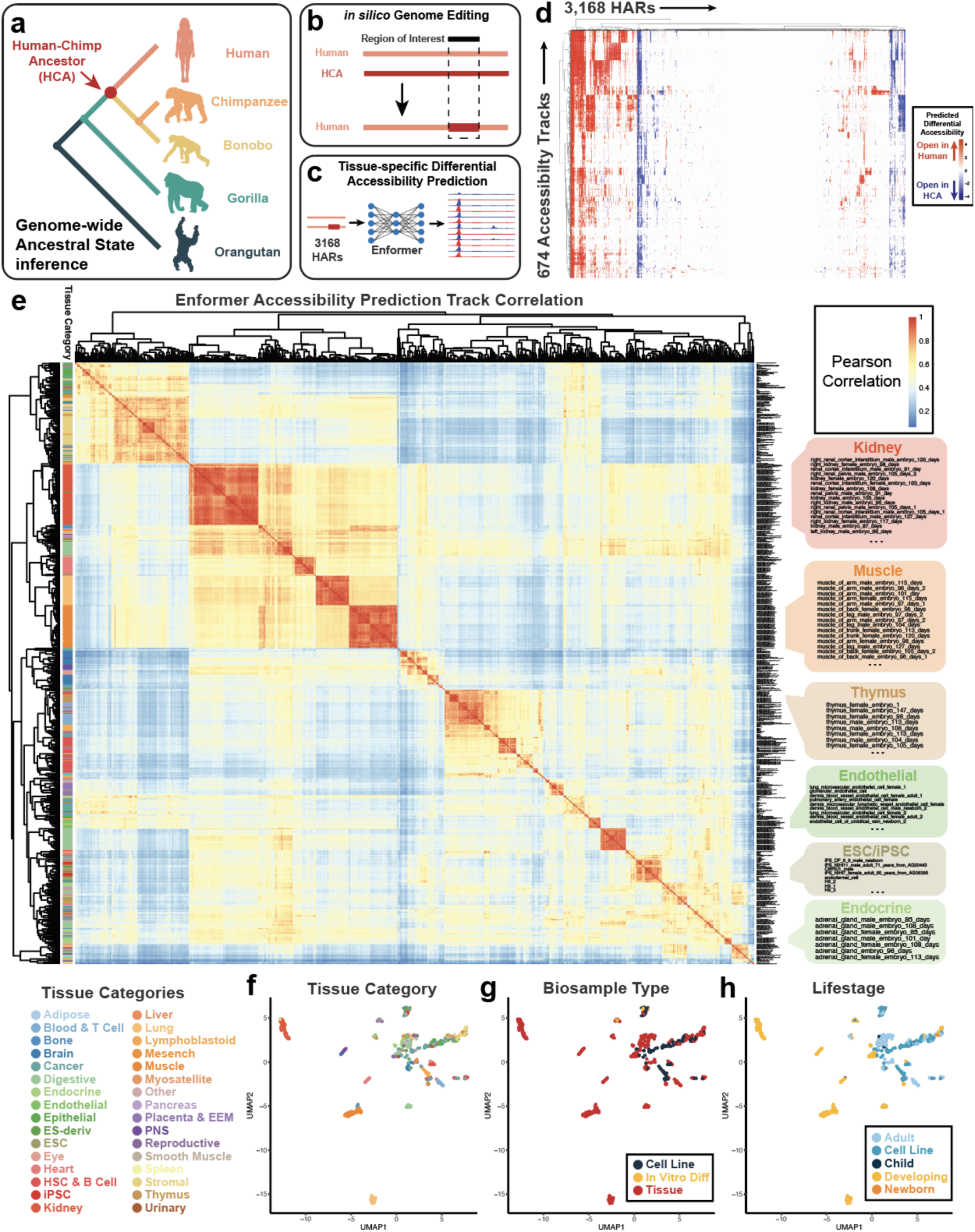
Correlation-based subsampling of Enformer accessibility prediction tracks. (**a**) Schematic of the human-chimpanzee ancestor (HCA) in the context of recent human evolution. (**b**) *In silico* genome editing of human accelerated regions (HARs), generating haplotypes containing the human reference sequence with ancestralized HAR sequences. (**c**) Haplotypes for modern humans and ancestralized haplotypes for 3,168 HARs were used to predict tissue-and cell-type-specific differential accessibility using Enformer. (**d**) Heatmap of predicted differential accessibility for 3,168 HARs across 674 biosamples. Red indicates regions with greater accessibility in humans while blue indicates greater accessibility in the HCA. (**e**) Correlation matrix of Enformer-predicted differential accessibility values across all 674 biosamples, clustered hierarchically. Tissue categories corresponding to clusters are highlighted on the right, including kidney, muscle, thymus, endothelial, ESC/iPSC, and endocrine tissues. (**f-h**) Uniform manifold approximation and projection (UMAP) of biosamples based on Enformer-predicted chromatin accessibility across HARs. Colors represent (**f**) 32 tissue categories (**g**) 3 biosample types (cell lines, tissues, or *in vitro* differentiated cells) or (**h**) 5 developmental stages (adult, cell line, child, developing, newborn).

**Extended Data Figure 2:**
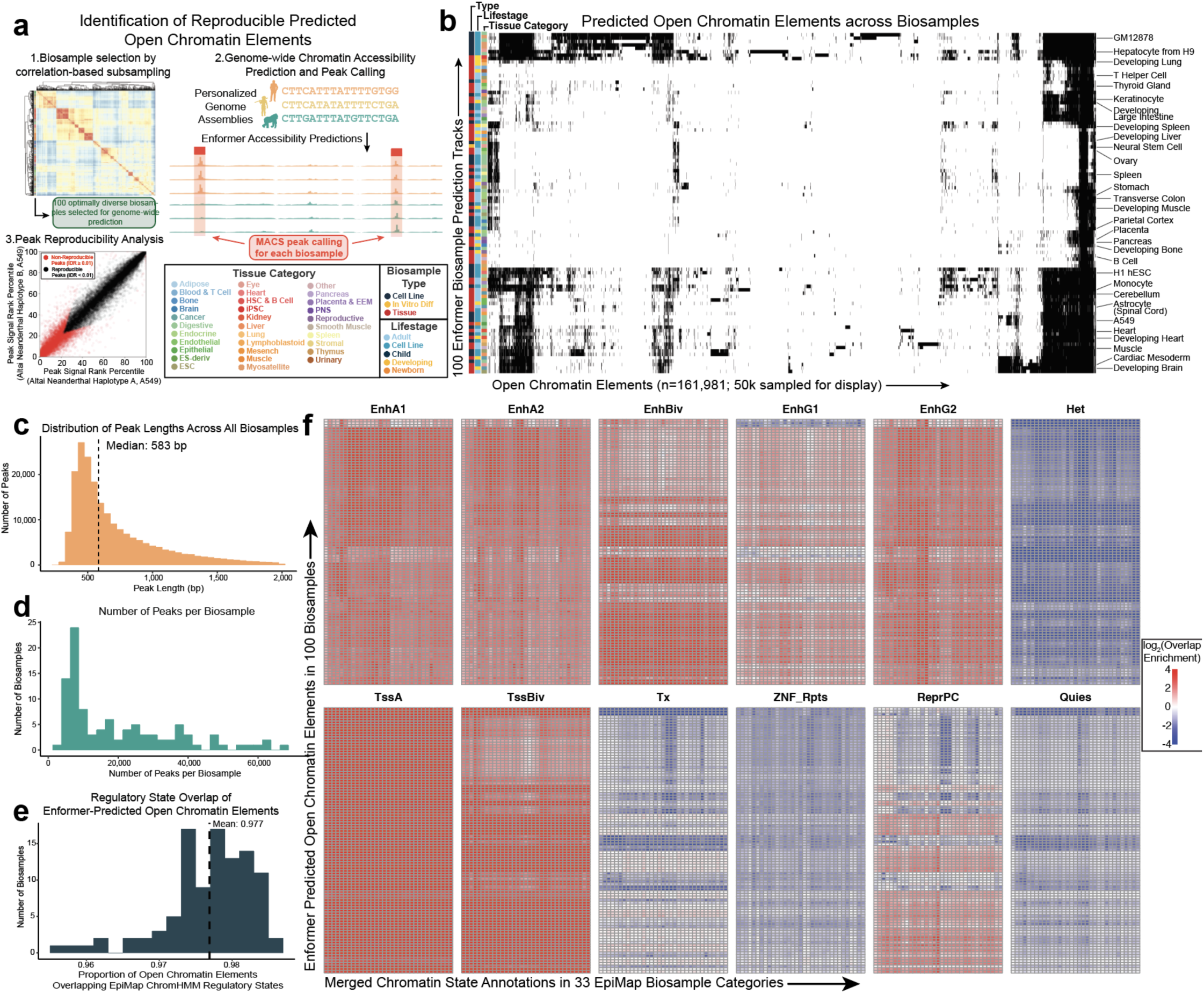
Identification and characterization of reproducible open chromatin elements from Enformer prediction tracks. (**a**) Workflow to identify reproducible open chromatin elements from Enformer-predicted chromatin accessibility tracks. To reduce computation, we selected 100 optimally diverse biosamples using correlation-based subsampling of Enformer chromatin accessibility prediction tracks. We predicted genome-wide accessibility using Enformer across 34 personalized genome assemblies and identified initial peak sets for each biosample using MACS3^1^. Final peak sets for each biosample were generated following IDR-based peak reproducibility thresholding. (**b**) Heatmap of predicted open chromatin elements (n=161,981, 50,000 sampled for display; columns) across 100 biosamples (rows). (**c**) Distribution of peak lengths for all peaks across biosamples. (**d**) Distribution of number of peaks per biosample. (**e**) Proportion of Enformer-predicted open chromatin elements overlapping regions annotated as active regulatory states (EnhA1, EnhA2, EnhBiv, EnhG1, EnhG2, EnhWk, TssA, TssBiv, TssFlnk, TssFlnkD, TssFlnkU) in at least one EpiMap biosample. (**f**) Heatmaps display log_2_-scaled overlap enrichments between Enformer-predicted open chromatin elements across 100 biosample contexts (rows) and chromatin state annotations from the EpiMap resource, aggregated across 33 biosample categories (columns). For each chromatin state, annotations were generated as the union of that state across all biosamples within each tissue category. Enformer-predicted open chromatin elements are strongly enriched for active enhancer (EnhA1, EnhA2, EnhBiv, EnhG1, EnhG2) and promoter (TssA, TssBiv) states and depleted from heterochromatin (Het), ZNF/repeats (ZNF_rpts), and quiescent (Quies) states.

**Extended Data Figure 3:**
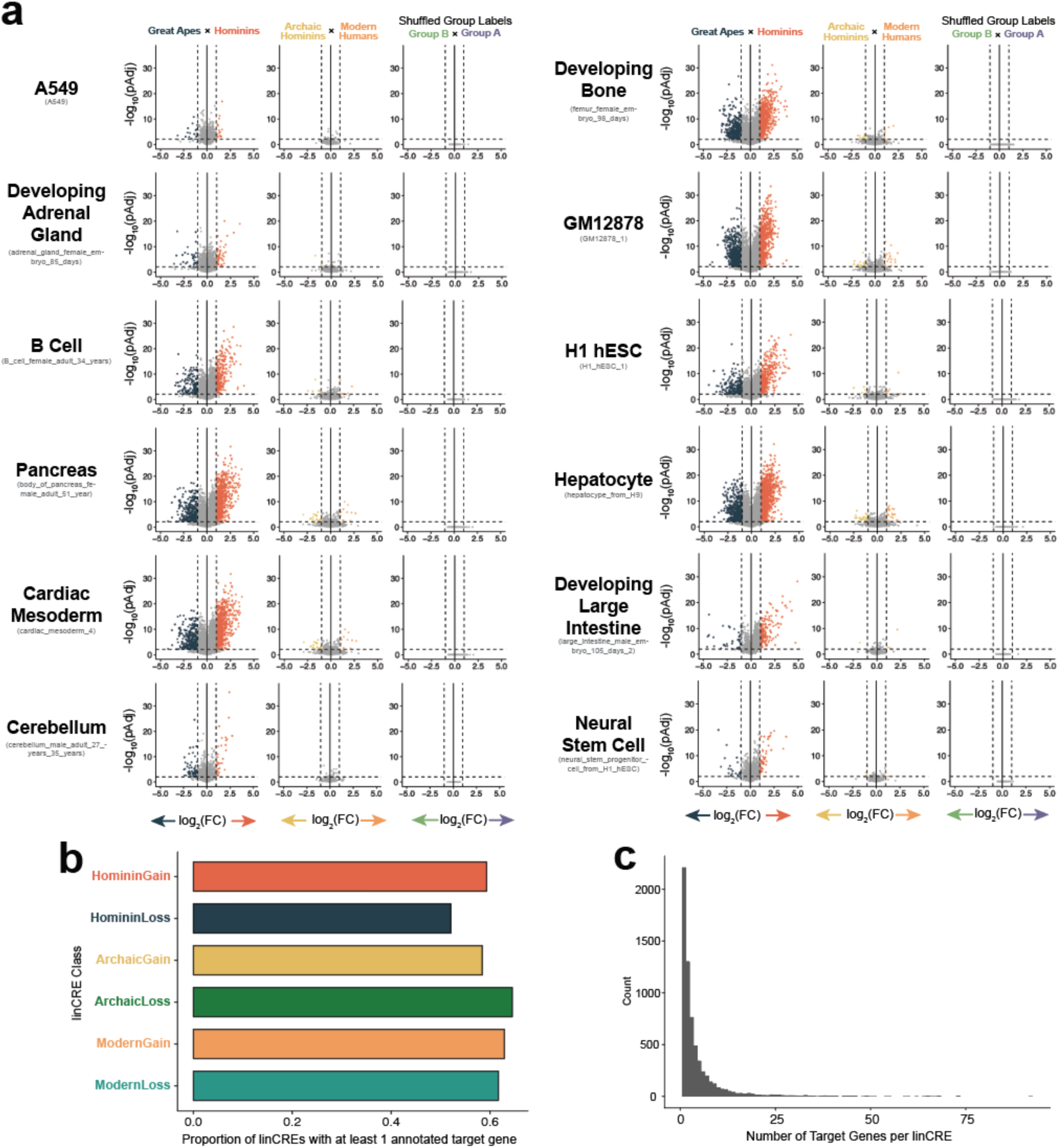
Extended visualization of differential accessibility analysis and target gene prediction. (**a**) Volcano plots display differential chromatin accessibility between hominins and great apes (left), modern humans and archaic hominins (middle), and shuffled group labels (right) in twelve representative biosamples. (**b**) Proportion of elements in each linCRE class with at least one annotated target gene in the EpiMap enhancer-gene linking resource^2^. (**c**) Distribution of the number of target genes per linCRE.

**Extended Data Figure 4:**
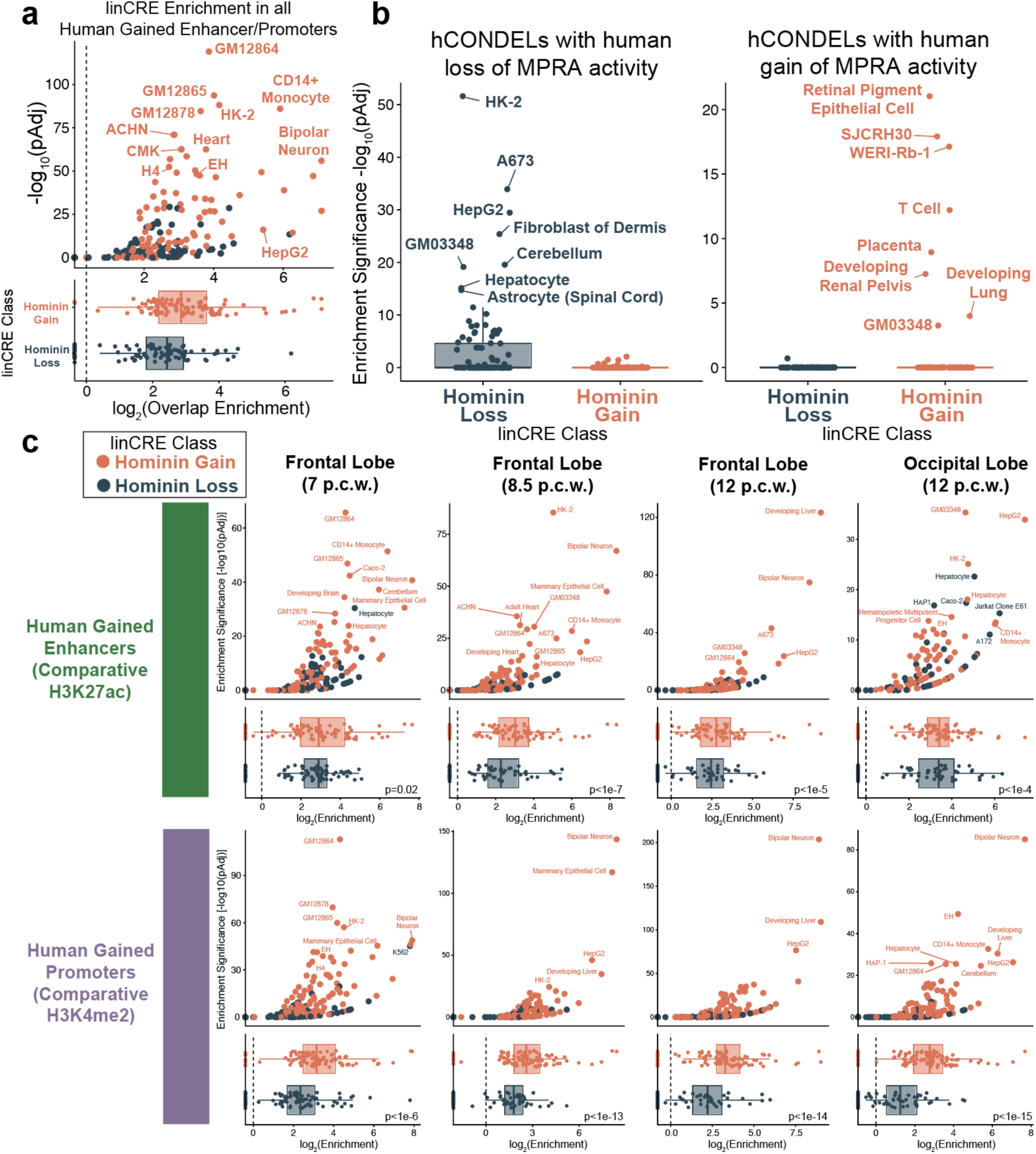
Predicted lineage-specific *cis*-regulatory elements are enriched for experimentally characterized human-ape regulatory differences. (**a**) (top) Volcano plot quantifies enrichments between linCREs, stratified by biosample, and human-gained enhancers/promoters (HGE/Ps) identified through cross-species ChIP-seq^3^. (bottom) Boxplots compare log_2_-scaled overlap enrichments between Hominin Gain-linCREs (orange) and Hominin Loss-linCREs (blue). (**b**) Overlap enrichment analysis between linCREs and human conserved deletions (hCONDELs) with either human loss of MPRA activity (left) or human gain of MPRA activity (right). (**c**) Extended overlap enrichments between predicted linCREs and human gained enhancers (top) and human gained promoters (bottom) stratified by time point and brain region of HGE/P ascertainment.

**Extended Data Figure 5:**
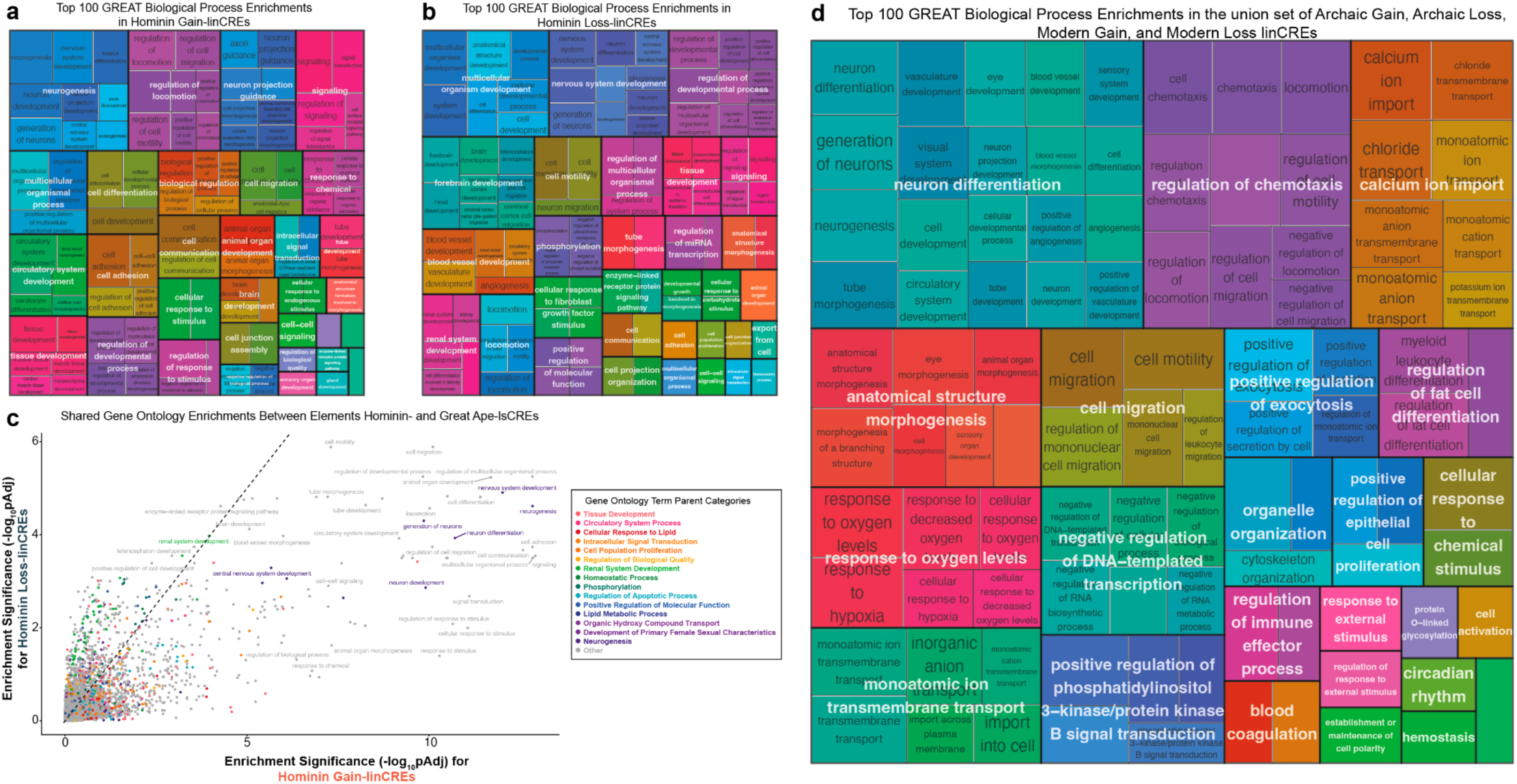
Extended visualization of biological process enrichment in linCREs. (**a-b**) The 100 most significant Gene Ontology (GO) biological process enrichments semantically clustered for visualization are displayed for (**a**) Hominin Gain-linCREs and (**b**) Hominin Loss-linCREs. Both sets are significantly enriched for neurogenic and developmental processes. (**c**) Shared gene ontology enrichments between Hominin Gain-and Hominin Loss-linCREs. Scatter plot displays enrichment significance (-log_10_-adjusted p-value) in Hominin Gain-linCREs (x-axis) and in Hominin Loss-linCREs (y-axis). Points are colored by their semantic cluster assignment, reflecting parent ontology categories. (**d**) Top 100 most significant GO biological process enrichment semantically clustered for visualization are displayed for the union set of Archaic Gain-, Archaic Loss-, Modern Gain-, and Modern Loss-linCREs.

**Extended Data Figure 6:**
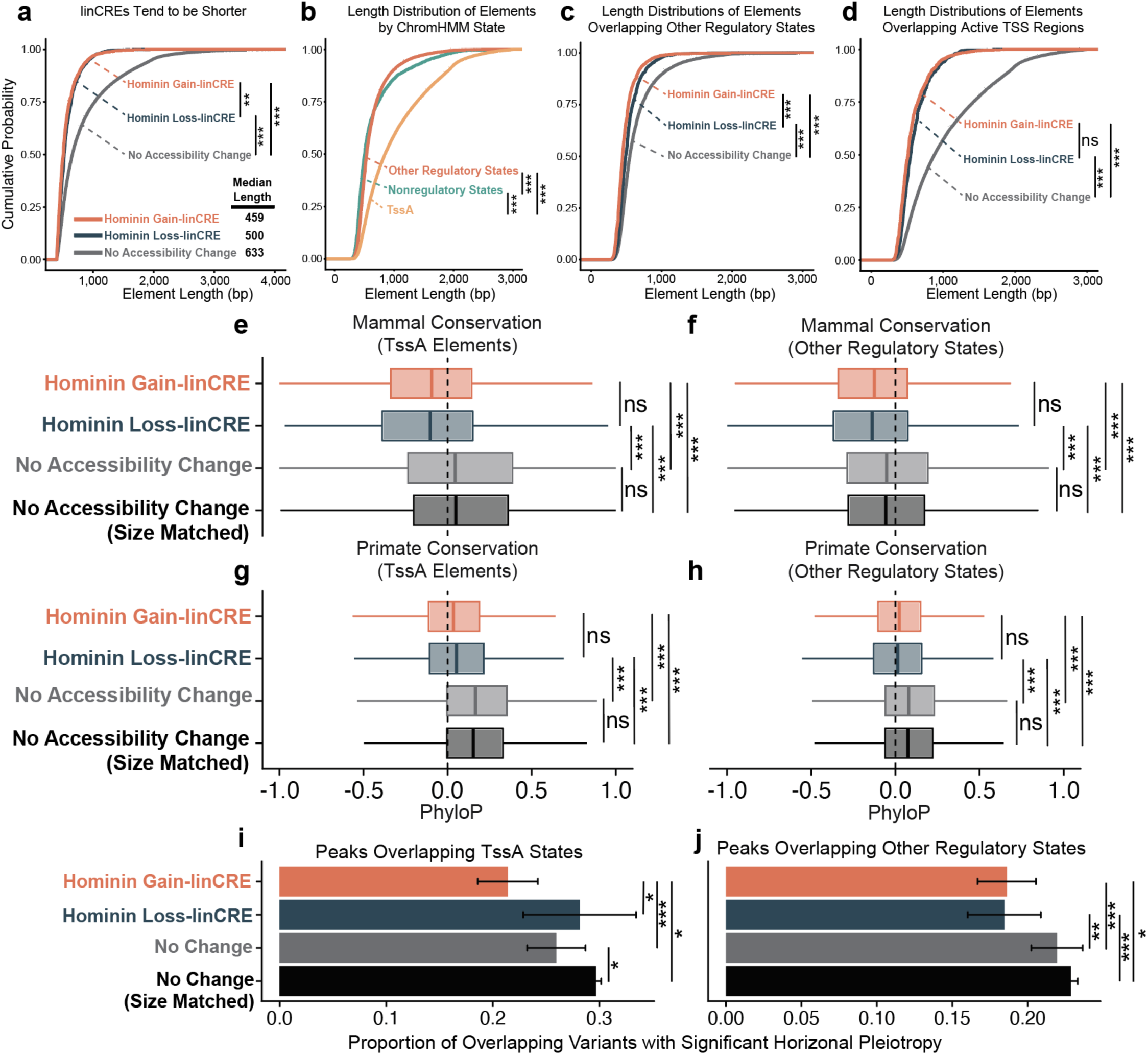
linCREs are shorter, less conserved, and less pleiotropic than elements with conserved accessibility. (**a**) Cumulative distribution of element lengths for peaks with no predicted accessibility change (gray), Hominin Gain-linCREs (orange), and Hominin Loss-linCREs (blue). (**b**) Element length distribution of open chromatin elements stratified by ChromHMM state. TssA elements (yellow) are significantly longer than elements in other regulatory states (orange) or elements outside of regulatory states (teal). (**c-d**) Length distributions for elements overlapping (**c**) non-TssA regulatory states or (**d**) TssA chromatin states. (**e-h**) PhyloP conservation scores for elements overlapping TssA (**e, g**) or other regulatory states (**f, h**), measured using mammalian (**e-f**) or primate (**g-h**) alignments. linCREs are less conserved, even when controlling for element length. (**i-j**) Proportion of overlapping variants with significant horizontal pleiotropy for elements overlapping TssA (**i**) or other regulatory ChromHMM states (**j**).

**Extended Data Figure 7:**
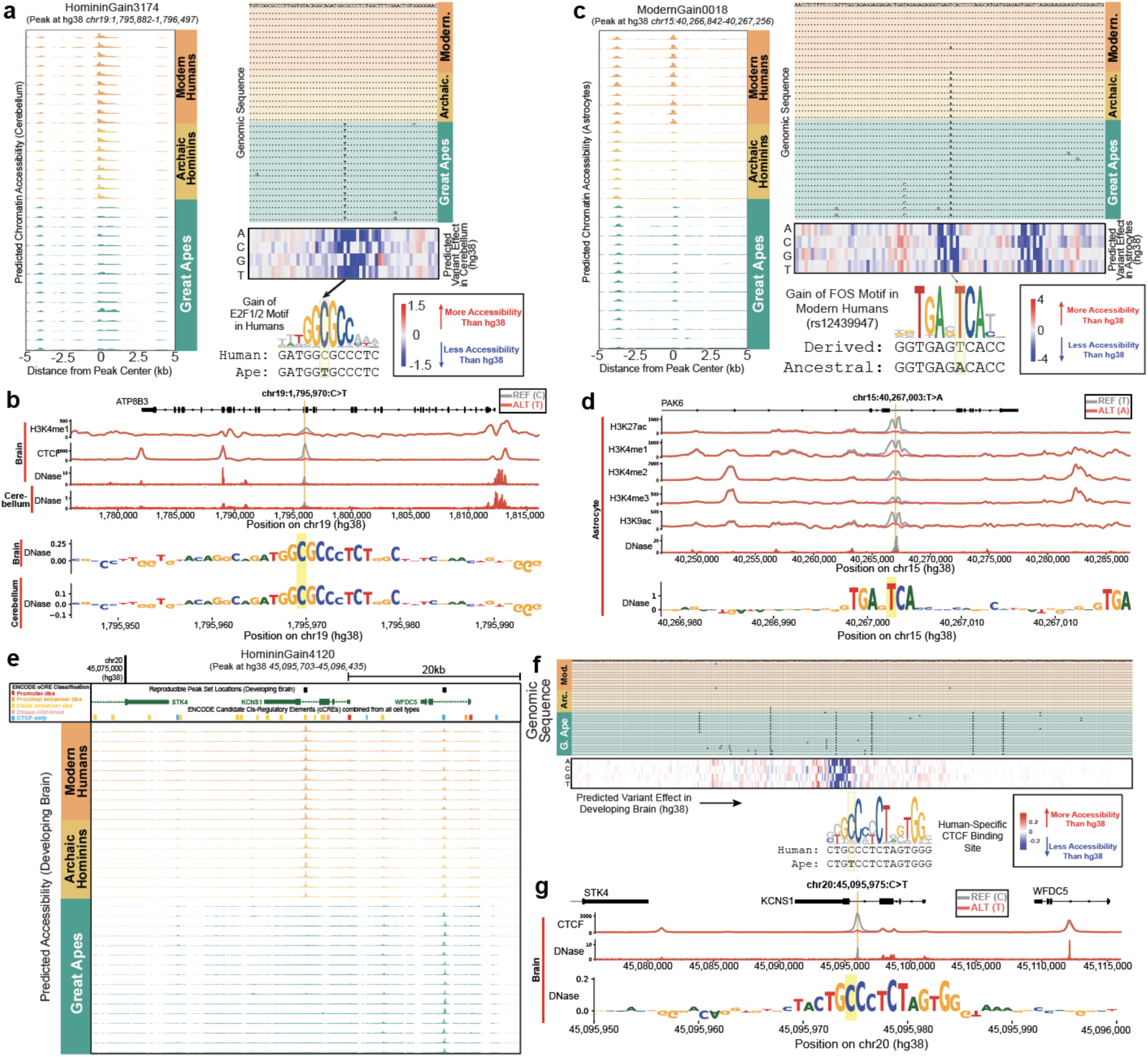
In silico saturation mutagenesis nominates single causal variants underlying lineage-specific regulatory activity. (**a**) Molecular dissection of a representative brain Hominin Gain-linCRE (HomininGain3174) linked^2^ to *GRIN3B*, a subunit of the N-Methyl-D-aspartic (NMDA) receptor implicated in schizophrenia and post-traumatic stress disorder^4,5^. Left, Enformer-predicted chromatin accessibility across all 34 haplotypes, demonstrating hominin-specific accessibility. Right, the genomic sequence surrounding a candidate SNP of interest (top); Enformer ISM showing that the derived hominin allele coincides with a base that strongly influences the accessibility prediction (middle); and the resulting gain of a novel E2F1/2 transcription factor binding site in hominins (bottom), providing a candidate molecular mechanism for the observed accessibility divergence. (**b**) Variant effect prediction for the prioritized variant in HomininGain3174 with AlphaGenome. Predicted brain H3K4me1, CTCF, and DNase profiles for the reference (C) and alternate (T) alleles, with cerebellum DNase sequence-contribution logos below. (**c**) Molecular dissection, as in (**a**), for a Modern Gain-linCRE (ModernGain0018) linked to *KNL1* in an astrocyte context. The derived modern human allele creates a FOS motif. This site is polymorphic in humans (rs12439947, MAF ∼0.17), and a single human haplotype carrying the ancestral allele shows markedly reduced predicted accessibility. (**d**) Variant effect prediction for the prioritized variant in ModernGain0018 with AlphaGenome, including predicted astrocyte H3K27ac, H3K4me1, H3K4me2, H3K4me3, H3K9ac, and DNase profiles for the reference (T) and alternate (A) alleles, with DNase sequence-contribution logos below. (**e**) Enformer-predicted chromatin accessibility across all 34 haplotypes (developing brain context) at a Hominin Gain-linCRE (HomininGain4120) in an intron of *KCNS1*, showing hominin-specific accessibility. (**f**) Sequence alignment, Enformer ISM matrix, and motif diagram for HomininGain4120, showing that a single derived allele creates a human-specific CTCF binding site predicted to drive accessibility in the developing brain. (**g**) Variant effect prediction for the prioritized variant in HomininGain4120 with AlphaGenome. Predicted brain CTCF and DNase profiles for the reference (C) and alternate (T) alleles, with DNase sequence-contribution logos below.

**Extended Data Figure 8:**
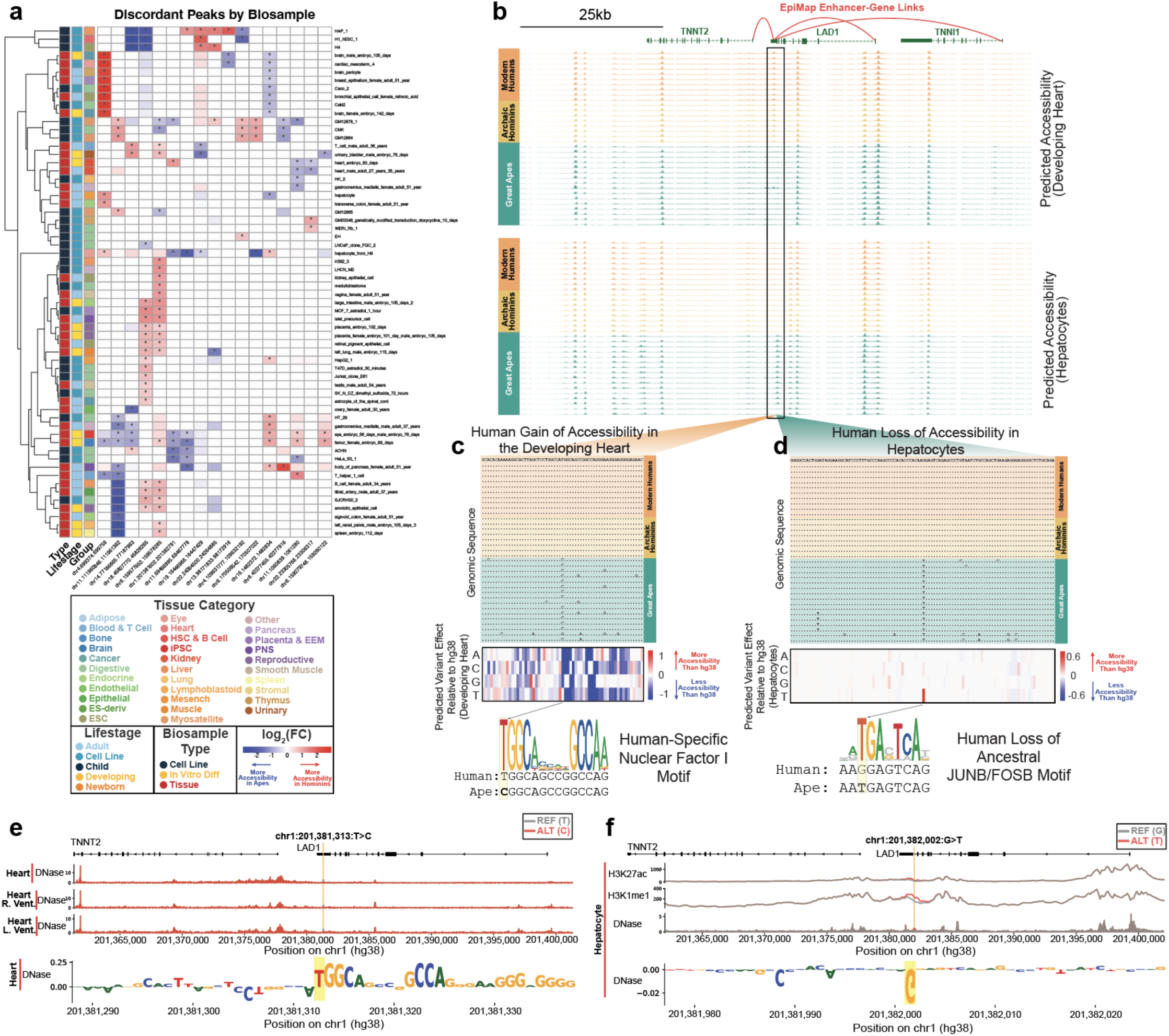
Complex regulatory innovation in a troponin-linked enhancer with tissue-discordant lineage specificity. (**a**) Heatmap displays differential accessibility patterns across 17 regulatory elements predicted to exhibit hominin-specific accessibility in at least one biosample context with predicted great ape-specific accessibility in another. Values represent log_2_ fold-change (hominin/ape) in predicted accessibility. (**b**) Predicted chromatin accessibility across haplotypes for a regulatory element in the 3’ UTR of *LAD1* with regulatory connections to *TNNT2* and *TNNI1*. This region harbors a Hominin Gain-linCRE (HomininGain0491) in the developing heart (top) that partially overlaps a Hominin Loss-linCRE (HomininLoss0259) in hepatocytes (bottom). (**c**) Saturation mutagenesis analysis in the developing heart identifies a high-impact SNP that generates a human-specific Nuclear Factor I motif. (**d**) Saturation mutagenesis in hepatocytes reveals a high-impact SNP corresponding to human-specific loss of an ancestral JUNB/FOSB motif. (**e**) Variant effect prediction for the prioritized variant in HomininGain0491 with AlphaGenome, supporting derived allele-specific accessibility in heart-related biosamples driven by the gain of a Nuclear Factor I motif. (**f**) Variant effect prediction for the prioritized variant in HomininLoss0259 with AlphaGenome, supporting ancestral allele-specific H3K27ac, H3K4me1, and DNase signals in hepatocytes.

**Extended Data Figure 9:**
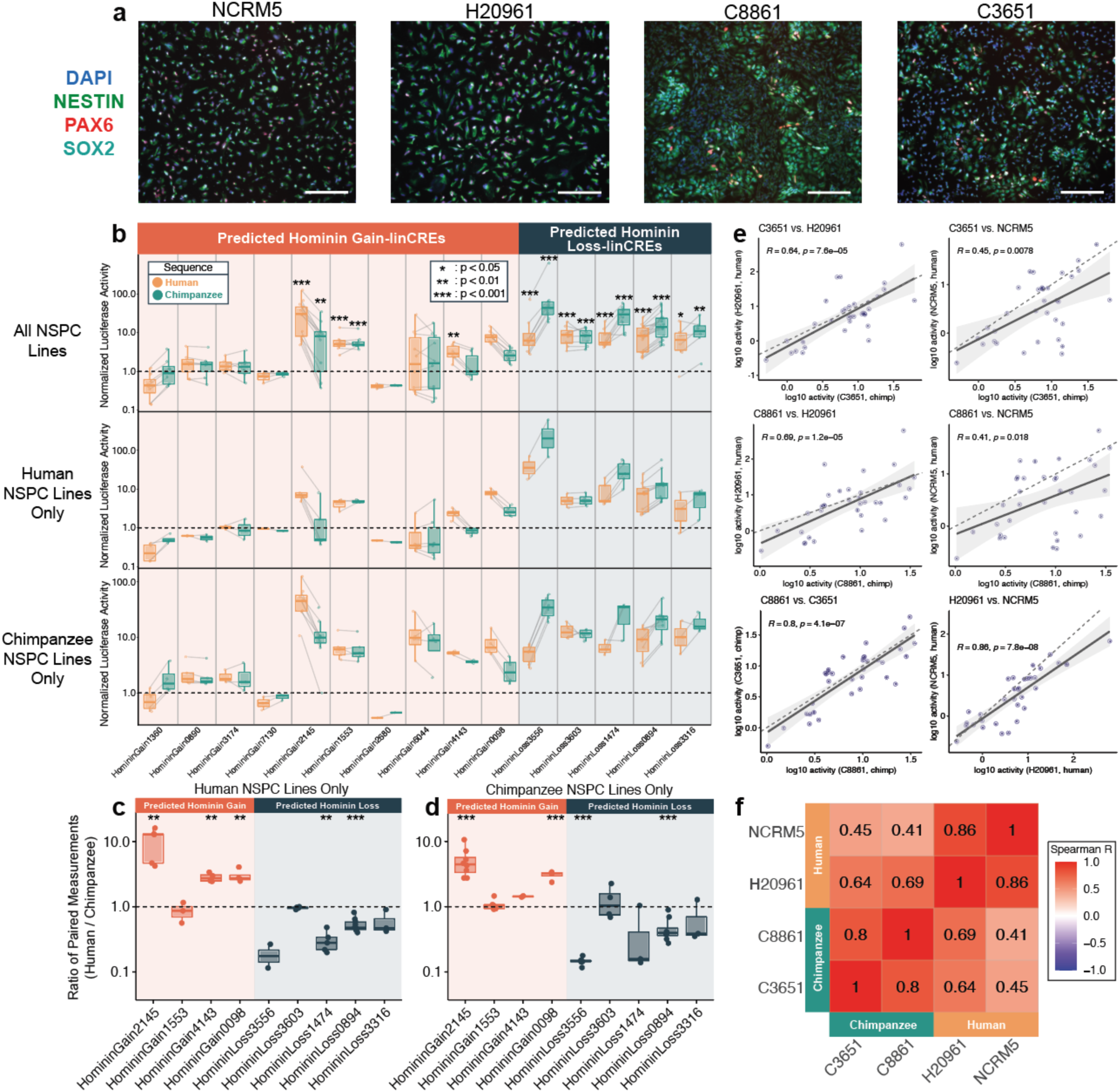
linCRE luciferase activity patterns persist across trans-regulatory environments. (**a**) Representative image of neural stem/progenitor cells (NSPCs) derived from four iPSC lines (human: H20961 and NCRM5. Chimpanzee: C8861 and C3651) used in luciferase reporter assays stained for DAPI, NESTIN, PAX6, and SOX2. Scale bar length = 170 μm. (**b**) Normalized luciferase for all 15 tested ortholog pairs (human, orange; chimpanzee, teal) grouped by predicted linCRE class (predicted Hominin Gain, left; predicted Hominin Loss, right). Data are shown across all NSPC lines (top), human NSPC lines only (middle), and chimpanzee NSPC lines only (bottom). Paired measurements are linked by gray lines. Asterisks in the top panel denote elements with significant enhancer activity above background (one-sided t-test, BH-adjusted P < 0.05). (**c-d**) Ratio of paired human/chimpanzee luciferase measurements for the nine active elements, displayed for data in (**c**) human NSPC lines only or (**d**) chimpanzee NSPC lines only. Direction and magnitude of species skew largely persists across trans-regulatory environments. Asterisks denote elements with significant enhancer activity above background (one-sided t-test, BH-adjusted P < 0.05). (**e**) Pairwise correlation of luciferase measurements for each construct across all six pairs of four NSPC lines. Luciferase activity for each construct was positively and significantly correlated across all pairs of cell lines (Spearman correlation ranging from 0.41 to 0.86). Within-species luciferase activity correlations (H20961 vs. NCRM5, R=0.86; C8861 vs. C3651, R=0.78) were higher than cross-species comparisons (R=0.41-0.69 across four comparisons), indicating modest trans-regulatory differences. Dashed line: y=x. Solid line and shaded band indicate trendline with 95% confidence interval. (**f**) Correlation matrix across cell lines. *: p < 0.05; **: p < 0.01; ***: p < 0.001.

**Extended Data Figure 10:**
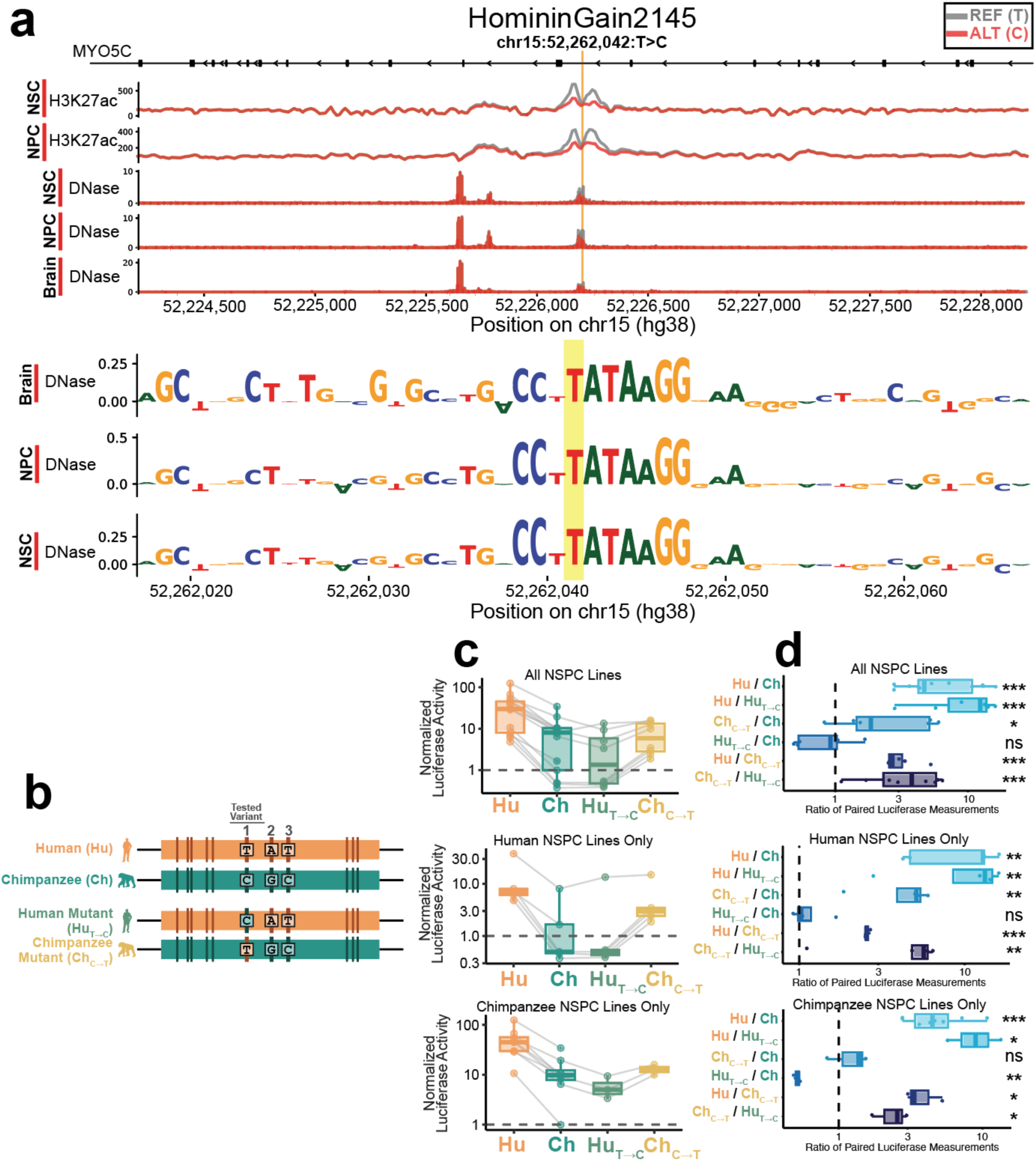
Extended analysis of HomininGain2145. (**a**) Variant effect prediction for the prioritized variant in HomininGain2145 with AlphaGenome, showing (top) Ref allele-specific H3K27ac signal in neural stem cell (NSC) and neural progenitor cell (NPC) contexts and Ref allele-specific DNase in NSCs, NPCs, and developing brain. (Bottom) DNase sequence contribution logos identify the prioritized variant within an SRF motif contributing to the accessibility prediction. (**b**) Luciferase construct design, reproduced from **Fig. 5c**. Human (Hu) and chimpanzee (Ch) reference sequences and reciprocal single-variant mutants at variant 1, including human-mutant (Hu_T→C_), the human sequence carrying the chimpanzee allele at variant 1 and chimpanzee-mutant (Ch_C→T_), the chimpanzee sequence carrying the human allele at variant 1. (**c**) Luciferase analysis of the ISM-prioritized causal variant in HomininGain2145, shown across (top) all NSPC lines, as in **Fig. 5c** (middle) human NSPC lines only and (bottom) chimpanzee NSPC lines only. (**d**) Ratio of paired luciferase measurements for construct comparisons shown across (top) all NSPC lines, as in **Fig. 5c** (middle) human NSPC lines only and (bottom) chimpanzee NSPC lines only. Asterisks denote elements with significant human-chimpanzee differences (t-test on log_2_ ratio, BH-adjusted). *: p < 0.05; **: p < 0.01.

## Methods

### Personalized genome assembly and multiple alignment

We generated unphased diploid genome assemblies for 17 individuals through reference-guided assembly. Our procedures for reference-guided assembly, benchmarking of reference-guided assembly through simulation, library selection, masking of problematic genomic regions, and construction of the 35-way multiple alignment are described in detail in **Supplementary Notes 1 and 2**.

### Correlation-based subsampling of Enformer prediction tracks

The Enformer model predicts chromatin accessibility from input genomic sequences for 674 distinct biosamples, encompassing a wide array of cell lines, primary tissues, and developmental timepoints. To reduce computational demands and disk space while ensuring broad representation of this phenotypic space, we performed correlation-based subsampling, selecting 100 of the 674 tracks for downstream analyses. This subsampling was based on Enformer-predicted differential accessibility in fast-evolving regions of the human genome, aligning our biosample selection with the study goal of identifying regulatory innovations in the human lineage.

We began by accessing a previously published genome-wide multiple sequence alignment of five primate species, including human (*hg38*), chimpanzee (*panTro6*), bonobo (*panPan2*), gorilla (*gorGor5*), and orangutan (*ponAbe3*) along with an inferred sequence of the human-chimpanzee ancestor (**Extended Data Fig. 1a**)^1^. To directly compare modern human and ancestral sequences, we developed the program *multiFaSequenceSwap*, as part of the Gonomics repository^2^, to perform “*in silico* genome editing”. In this process, we substitute a genomic region of interest - defined by a BED file - with the orthologous sequence from another species using a FASTA format multiple alignment (**Extended Data Fig. 1b**). We applied this program to “ancestralize” 3,168 human accelerated regions (HARs) - genomic regions with elevated rates of human-specific divergence despite strong cross-species conservation across non-humans species^3^. Critically, by reverting only mutations within HARs to their ancestral states while preserving the surrounding sequence, we were able to isolate the specific effects of these mutations on chromatin accessibility. HAR locations were accessed from the Gene Expression Omnibus (GEO) at accession number GSE180714. Next, we used Enformer^4^ to predict chromatin accessibility at each HAR for both the human reference sequence and the ancestralized sequence (**Extended Data Fig. 1c**).

Enformer predicts chromatin accessibility at a resolution of 128bp windows. In this analysis, we calculated chromatin accessibility as the maximum amplitude in a region of 7 windows (896bp) with the window overlapping the center of the queried region as the central window.

We then calculated the log_2_-scaled differential accessibility between Enformer-predicted chromatin accessibility in the human and chimpanzee haplotype (**Extended Data Fig. 1d**). To prioritize biologically meaningful accessibility differences in regulatory elements over potential technical noise, log_2_-scaled fold changes were masked to 0 if the difference between the human and ancestralized sequence was below 0.5. Additionally, we truncated absolute log_2_-scaled fold changes greater than 5 to a maximum of 5 to control for extreme outliers that could disproportionately influence downstream clustering. Finally, we calculated a biosample-biosample correlation matrix from the resulting vectors of log_2_-scaled fold change across all HARs (**Extended Data Fig. 1e**) and used hierarchical clustering to select 100 optimally diverse biosamples.

We visualized regulatory relationships between biosamples using a uniform manifold approximation and projection (UMAP) using the R UMAP package^5^.

For clarity and readability in figures, we replace original biosample names with shortened, human-readable display names by standardizing punctuation and removing metadata. For example, ‘T_cell_male_adult_36_years’ is displayed as ‘T Cell’, and ‘left_lung_male_embryo_115_days’ is presented as ‘Developing Lung’.

### Genome-wide chromatin accessibility prediction with Enformer

We developed a computational pipeline to perform genome-wide chromatin accessibility prediction using the pre-trained Enformer model. This pipeline requires several input files. First, it accepts a query genome in FASTA format, along with supporting files. In this study, we used ANCoRA-derived genome assemblies and generated the corresponding index (.fai format) and chromosome sizes (.chrom.sizes format) files using *samtools faidx* and the kentUtil command *faSize -detailed*, respectively^6,7^. Second, the pipeline accepts a tab-separated table of prediction track names and indices, corresponding to the 100 diverse biosamples identified in the Method section “Correlation-based subsampling of Enformer prediction tracks”. Third, it requires a directory path to the pre-trained Enformer model.

Using these inputs, our pipeline scans the genome in non-overlapping sliding windows, with each window size matched to the Enformer context window. Chromatin accessibility predictions for each 128 bp prediction window are generated and written to output files in the UCSC bedGraph format. These files are subsequently converted to bigWig format using the Gonomics tool *bedGraphToWig* and the kentUtil command *wigToBigWig*^2,7^.

While the current analysis was restricted to Enformer-based chromatin accessibility predictions, this genome scanning framework can be extended to other Enformer prediction tracks, which include diverse Chromatin Immunoprecipitation (ChIP)-seq and Cap Analysis of Gene Expression (CAGE) tracks from diverse biosamples.

Predictions were generated in strictly contiguous, non-overlapping 114,688bp prediction windows tiled from the 5’ end of each chromosome. To generate predictions for each 114,688 bp target interval, we extracted a 393,216 bp sequence centered on the target interval to provide the flanking context required by the model’s receptive field, padding with ’N’s where the context window extended beyond chromosome boundaries. Notably, the terminal 3’ remainder of each chromosome (shorter than one full prediction window) was not predicted. Across personalized assemblies, unpredicted regions accounted for approximately 0.04% of assembly bases.

### Defining open chromatin elements

We developed a custom pipeline to identify open chromatin elements from Enformer-derived chromatin accessibility prediction tracks that includes peak calling and reproducibility analysis to limit our analysis to high-quality peaks.

Initial peak sets were called from each bigWig format accessibility prediction track with MACS3 bdgpeakcall^8^. To determine the optimal significance threshold for peak calling, we used the MACS3 cutoff analysis tool across all biosamples and haplotypes, which determines the number of peaks as a function of the significance threshold score. We defined an adaptive threshold at the elbow point of this curve for each prediction track, ensuring consistent peak quality across diverse biosamples and haplotypes.

As raw peaks were called on haplotype-specific assembly coordinates, we used the UCSC liftOver tool to map peak coordinates to *hg38* coordinates. Peaks that failed to map or mapped to non-canonical chromosomes were discarded.

To generate a final set of high confidence peaks for each biosample track, we assessed the reproducibility of peaks within assembly groups (modern humans, archaic hominins, and great apes) by applying the Irreproducible Discovery Rate (IDR) framework developed and applied in the ENCODE and modENCODE consortia^9–11^. Briefly, the IDR statistic evaluates the consistency of peaks between pairs of samples, where lower IDR scores indicate higher reproducibility. For each biosample, we defined a peak as reproducible within a group if IDR < 0.01 in at least one pairwise comparison among haplotypes within that group. Finally, we defined a final peak call set for each biosample as the union of reproducible peaks across the three groups.

### Chromatin State Annotation Enrichments

To evaluate whether Enformer-predicted open chromatin elements correspond to known regulatory elements, we performed overlap enrichment analyses against chromatin state annotations from the EpiMap resource^12^. To this end, we accessed ChromHMM-derived chromatin state annotations across 833 biosamples, categorized into 33 tissue groups. For each chromatin state, we generated a union set by merging all occurrences of that state across all biosamples within each EpiMap tissue group. We evaluated overlap enrichments between EpiMap tissue-specific chromatin state annotations and Enformer-derived reproducible predicted open chromatin elements using the Binomial-based *overlapEnrichments* tool from Gonomics^2^. Resulting enrichment scores were log-transformed and visualized with pheatmap.

### Differential accessibility analysis

To identify differential accessibility events across haplotypes, we first quantified predicted chromatin accessibility for each haplotype across the set of reproducible open chromatin elements identified for each biosample. As reproducible peaks were defined on hg38 reference coordinates, we used the UCSC liftOver tool to convert each peak set to personalized genome coordinates. Notably, to address positional instability of open chromatin elements across haplotypes, we generated separate peak sets for each biosample. This approach ensured that differential accessibility was assessed based on peak positions specific to each biosample and avoided the merging of partially overlapping peaks at superenhancers, where individual constituent enhancers may not be active in a given biosample. Next, we used the kentUtil *bigWigAverageOverBed* to quantify predicted chromatin accessibility for each element, calculating the average predicted DHS-seq signal across the length of each peak. For each biosample, we reorganized these results into matrices reporting the mean accessibility prediction for each peak across all haplotypes.

We used these matrices to test each open chromatin element for differential accessibility in two comparisons: (1) between modern humans and archaic hominins, and (2) between hominins (encompassing both modern humans and archaic hominins) and great apes. To ensure robustness, we also conducted differential accessibility analysis on peaks after shuffling group labels. Open chromatin elements were defined as differentially accessible at a threshold of an FDR-adjusted p value < 0.01 (t test) and an absolute log2 fold change > 1. We also conducted this analysis after permuting group labels.

We named each class of lineage-specific *cis*-regulatory elements (linCREs) by the lineage and direction of its accessibility change relative to the inferred ancestral state, using great apes as the outgroup. Hominin-great ape events were classified as Hominin Gain (elevated accessibility in hominins) or Hominin Loss (elevated accessibility across great ape species, interpreted as loss of ancestral activity in the human lineage).

To categorize linCREs from the comparison between modern humans and archaic hominins, we polarized each differentially accessible region against the great ape ancestral state by assessing whether it overlapped a reproducible open chromatin element in the corresponding great ape biosample. Differentially accessible regions with elevated accessibility in modern humans that lacked an overlapping great ape peak were classified as Modern Gain, whereas those overlapping a great ape peak were classified as Archaic Loss, consistent with loss of ancestral regulatory activity in the archaic lineage. Conversely, regions with elevated accessibility in archaic hominins were classified as Archaic Gain when no overlapping great ape peak was present and as Modern Loss when an overlapping great ape peak indicated ancestral accessibility.

### Enhancer-gene target prediction

To link predicted linCREs to putative target genes, we utilized tissue-specific enhancer-gene linking predictions from the EpiMap resource^12^. We accessed raw enhancer-gene links for all available EpiMp tissue groups and parsed these files into BED format, mapping Ensembl gene IDs from the EpiiMap datasets to their corresponding gene symbols using GENCODE v44 gene annotations^13^. Because EpiMap annotations were available on hg19 coordinates, we converted enhancer coordinates to the hg38 assembly using the UCSC liftOver tool^7^. We then used the Gonomics intervalOverlap command to intersect our predicted linCREs with EpiMap enhancer coordinates. We provide a complete set of putative target genes for each linCRE, defined by aggregating all overlapping enhancer-gene links across all EpiMap tissue groups, as **Supplemental Table 8**.

### Overlap Enrichment Comparisons to Previously Described Regions of Evolutionary Interest

To validate the biological relevance of Enformer-predicted linCREs, we assessed overlap enrichments between these elements and experimentally characterized sets of human-ape regulatory differences, sequence-based annotations of evolutionarily significant regions on the human lineage, and human-ape differentially expressed genes.

We compared 200 sets of predicted linCREs (100 biosamples for both Hominin Gain and Hominin Loss linCREs) to human conserved deletions (hCONDELs) tested for differential regulatory activity through massively parallel reporter assays (MPRAs)^14^. hCONDELs are genomic elements that are conserved across nonhumans but absent in the human genome due to recent deletions^15^. To this end, we accessed Table S1 of Xue et al, which contains processed DESeq2 results from an MPRA across 10,032 hCONDEL elements conducted across six cell lines: K562, GM12878, HEK293, NPC, SK-N-SH, and HepG2 cells. From these data, we identified hCONDELs with significant species-specific activity as those with a species skew BH adjusted P value < 0.05 and significant enhancer activity in at least one species (BH adjusted P < 0.1 from either the human or macaque haplotype). For each cell type, we partitioned the set of species-specific elements into those with human gain of MPRA activity (log_2_FC > 0) and those with human loss of MPRA activity (log_2_FC < 0). We then defined the set of all hCONDELs with human gain of MPRA activity (n=422) as the union set of hCONDELs with human gain of MPRA activity across all six cell types, which we generated with the Gonomics *bedMerge* program^2^. Similarly, we defined the set of all hCONDELs with human loss of MPRA activity (n=422) as the union set of hCONDELs with human loss of MPRA activity across all six cell types.

We compared Enformer-predicted DA regions to human-gained enhancers (HGEs) and human-gained promoters (HGPs) identified by Reilly et al^16^. HGE/Ps were identified using comparative ChIP-seq (H3K27ac for enhancers; H3K4me2 for promoters) across humans, rhesus macaques, and mice, focusing on four developing brain regions: frontal lobe at 7 post-conception weeks (p.c.w), 8.5 p.c.w., and 12 p.c.w., and occipital lobe at 12 p.c.w. We accessed all 8 sets of HGE/Ps(H3K27ac and H3K4me2 for all four brain samples) from the Gene Expression Omnibus (GSE63649) and converted genomic coordinates from *hg19* to *hg38* using the UCSC *liftOver* tool^7^. In addition to these 8 sets, we generated a union set of all HGE/Ps (n=13,390) with the Gonomics *bedMerge* program^2^.

For both hCONDELs and HGE/Ps, we evaluated overlap enrichments against Enformer-predicted linCREs with the Gonomics *overlapEnrichments* program, which performs a binomial-based enrichment test comparing observed to expected overlaps under a random distribution assumption in a defined genomic background. Given that predicted linCREs are strongly enriched in regulatory elements (**Extended Data Fig. 2f**), we defined the genomic search space for enrichment analysis as the complete set of open chromatin elements evaluated for differential accessibility in each biosample. This ensures that enrichments for predicted DA events are interpreted in the context of open chromatin elements rather than the entire genome, similar to the background region strategy employed by the GREAT tool. Enrichment significance is reported as the minus log_2_-scaled FDR-adjusted p value.

We accessed hg38 coordinates of UNICORNs and human ancestor quickly evolved regions (HAQERs) from the supplementary materials of their respective publications^1,17^. We accessed a set of HARs in hg19 coordinates used by Doan and colleagues^18^, comprising 2,737 regions aggregated from several earlier screens^19–24^. After liftover to hg38, 2,734 HARs were retained for analysis. For set level enrichment analyses not stratified by biosample, we used the Gonomics *overlapEnrichments* program using the union set of all open chromatin elements across all biosamples as background regions.

### Differentially Expressed Gene Overlap Enrichment Analysis

We assessed whether predicted linCREs are enriched near genes differentially expressed between humans and great apes. We obtained differentially expressed genes (DEGs) from Jorstad et al, who performed single-cell RNA sequencing of the middle temporal gyrus in adult humans, chimpanzees, and gorillas^25^. We mapped DEG gene IDs to GENCODE v44 gene annotations (hg38) to identify canonical transcription start site (TSS) coordinates^13^. We retained DEGs with an adjusted p-value below 0.001 and a log2-scaled fold change above 2, and split these gene sets by direction of differential expression (upregulated or downregulated in humans). To evaluate proximity of divergent distal enhancers to DEG loci, we extended each TSS ± 50 kb using the Gonomics tool *bedFormat*, and collapsed overlapping regions using *bedMerge*^2^. Enrichment of overlap between Enformer-predicted elements and these regions was quantified using the *overlapEnrichments* method as described above.

### Evaluating Functional and Regulatory Features of Predicted Divergent Elements

#### Chromatin State Annotations

To quantify the functional significance of open chromatin regions across evolutionary categories (Hominin Gain-linCRE, Hominin Loss-linCRE, and No Accessibility Change), we calculated the fraction of elements overlapping EpiMap-defined active regulatory states^12^. We generated two union sets of annotations across all 833 biosamples. The first contained all regions annotated as active promoters (TssA). The second included all regulatory states: EnhA1, EnhA2, EnhBiv, EnhG1, EnhG2, EnhWk, TssA, TssBiv, TssFlnk, TssFlnkD, and TssFlnkU.

Each set of open chromatin elements was intersected with these annotation sets using the Gonomics *intervalOverlap* tool^2^ to determine the proportion of elements overlapping active promoter regions, other regulatory annotations, or no regulatory annotations. To test for significant differences in regulatory content between groups, we performed pairwise Fisher’s exact tests using 2x2 contingency tables for TssA vs. non-TssA annotations and regulatory vs. non-regulatory annotations.

#### Gene Ontology Enrichment Analysis

We performed gene ontology (GO) enrichment analysis on Enformer-predicted linCREs to identify associations with biological process, cellular component, and molecular function ontologies using rGREAT, an R implementation of the Genomic Regions Enrichment of Annotations Tool^26,27^. We used the set of all open chromatin elements analyzed in this study as background regions to ensure that enrichments reflected functions specific to the differentially accessible subset of open chromatin elements, rather than broader patterns of regulatory activity inherent in open chromatin elements. To visualize and interpret enrichment results, we used the R package rrvgo to calculate semantic similarities between GO terms, reducing redundant terms into higher-order ontological themes and to visualize enrichments as tree maps^28^.

### Length Distribution Analysis and Stratified Sampling

To investigate the relationship between open chromatin element length and divergent accessibility, we compared length distributions across categories using Bonferroni-corrected pairwise t tests and visualized the cumulative density functions for each set of open chromatin elements.

To control for potential biases in select downstream analyses, we generated a size-matched set of ‘No Accessibility Change’ elements using a quantile binning approach. Specifically, we used the empirical length distribution of divergent elements (including both Hominin Gain-and Hominin Loss-linCREs) to define bin intervals. ‘No Accessibility Change’ elements were then partitioned into these bins, and from each bin, we randomly subsampled a number of elements matching the bin-wise count of divergent elements.

### Sequence Conservation Analysis

To evaluate selective constraint across CREs, we accessed two complementary genome-wide sequence conservation datasets in UCSC BigWig format, both providing single-base pair resolution conservation annotations. These datasets present phyloP scores, which quantify the deviation of observed substitution rates from neutral expectations^29^. Positive PhyloP scores indicate sequence conservation, reflecting selective constraint across macroevolutionary timescales and suggesting putative functional importance. In contrast, PhyloP scores near zero reflect neutral evolution, and negative scores indicate accelerated divergence.

We first accessed “Primate PhyloP” scores derived from a 239-way multiple sequence alignment of primates^30^. Additionally, we accessed “Mammal PhyloP” annotations derived from a multiple alignment of 447 diverse mammal genomes, combining the 241-way mammal alignment generated by the Zoonomia consortium^17^ with the additional primate genomes from Kuderna and colleagues.

We calculated the mean primate and mammal phyloP score for each Enformer-predicted open chromatin element using the *bigWigAverageOverBed* tool from the UCSC Genome Browser^7^. We compared the distributions of mean phyloP scores across two distinct partitionings of open chromatin elements. We conducted Bonferroni-corrected pairwise t tests to quantify differences in sequence conservation across linCRE classes. Nonsignificant p values are reported as “ns” above a threshold of p=0.01 in figure panels.

### Horizontal Pleiotropy Score Quantification

To assess the degree of horizontal pleiotropy associated with CREs, we accessed a dataset of 1,183,386 human genetic variants annotated with linkage disequilibrium (LD)-corrected horizontal number of traits pleiotropy scores (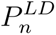 generated by Jordan et al.^31^). Briefly, 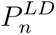 scores represent the expected value of the number of statistically independent traits for which a given variant is associated in a set of 100 traits. Each variant is annotated with both the 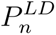 score as well as a p-value, reflecting the statistical significance of horizontal pleiotropy for the variant.

We lifted the original hg19 annotations to hg38 coordinates using the UCSC *liftOver* tool, retaining 1,182,918 genetic variants. We used the Gonomics tool *intervalOverlap* to identify variants overlapping Enformer-predicted open chromatin elements. We partitioned overlapping variants into those overlapping open chromatin elements (1) Hominin Gain-linCREs (2) Hominin Loss-linCREs or (3) CREs with no predicted change in accessibility.

For each set of genomic regions, we quantified the proportion of overlapping variants with significant horizontal pleiotropy, using a p-value threshold of 0.001. To visualize the uncertainty associated with these proportion estimates, we calculated 95% confidence intervals using Wald’s method. Finally, we conducted a Chi-square test to evaluate statistically significant differences in horizontal pleiotropy between sets of genomic elements.

### Sequence Divergence Analysis

To quantify the number of human-ape sequence differences within open chromatin elements, we used a previously generated set of variant calls representing base-level differences between the human reference genome (hg38) and the inferred human-chimpanzee ancestor sequence^1^. These differences were identified from a five-way whole-genome alignment of great apes and include both single nucleotide variants and short insertions or deletions. We used the Gonomics *intervalOverlap* command to intersect these variant calls with open chromatin elements to count the number of sequence differences overlapping each element. Elements were annotated with the total number of overlapping differences, which we used as a proxy for sequence divergence.

To compare sequence divergence across functionally divergent and conserved regulatory elements, we computed the number of sequence divergence per 100 base pairs and visualized cumulative distribution functions of divergence for each element class (*Hominin Gain-linCRE*, *Hominin Loss-linCRE*, *No Accessibility Change*, and *No Accessibility Change [Size Matched]*). Sequence divergence distributions were compared using Bonferroni-corrected pairwise *t*-tests.

### Accessibility Breadth Analysis

To quantify the regulatory breadth of open chromatin elements, we determined the number of distinct biosamples in which each element was predicted to be accessible. We compared 4 sets of genomic elements, including Hominin Gain-linCREs, Hominin Loss-linCREs, and the union set of Modern/Archaic-linCREs (Archaic Gain, Archaic Loss, Modern Gain, Modern Loss) and the size-matched set of CREs with conserved accessibility. We aggregated the four classes of Archaic/Modern-linCREs to achieve a set size supporting statistical comparisons. The size-matched conserved control was constructed to match the length distribution of Hominin Gain-and Hominin Loss-linCREs (see *Length Distribution Analysis and Stratified Sampling*). This control was also used as the comparator for the pooled Archaic/Modern-linCRe set, which was not itself used to define the length-matching bins. For each set, we concatenated, sorted, and merged overlapping intervals across all biosample-specific prediction tracks. For each merged genomic region, we counted the number of distinct contributing biosamples to define its accessibility breadth.

To statistically compare accessibility breadth across sets, we calculated the proportion of elements accessible in >15 biosamples. We estimated 95% confidence intervals for these proportions using Wald’s method, bounding the limits to [0,1]. We evaluated pairwise differences in accessibility breadth (>15 versus ≤ 15) between elements sets using Fisher’s exact tests, with p-values adjusted using the Benjamini-Hochberg false discovery rate procedure.

### In silico saturation mutagenesis and variant effect prediction

To prioritize causal variants within linCREs, we performed in silico saturation mutagenesis (ISM) using Enformer. For each element,we extracted the reference sequence in the model input window and substituted each of the three alternative nucleotides at every position, recording the predicted change in accessibility for the relevant epigenomic track relative to the reference allele, using either the human reference genome hg38 or the chimpanzee reference genome panTro6, as indicated in figures.

To assess robustness to model choice, we re-evaluated variants using AlphaGenome, a successor model to Enformer released after our initial screen^32^. For allele-specific epigenomic track visualizations, we extracted a 1,048,576bp sequence context centered on the variant from the hg38 reference genome. We generated predictions for both the reference and alternate alleles and visualized relevant epigenomic outputs subsetted to biologically relevant cell and tissue ontologies. To perform AlphaGenome-based sequence contribution logos, we performed localized *in silico* saturation mutagenesis. For this step, we utlized a 16kb sequence context window and exhaustively mutated every base across a 256bp interval centered on the prioritized variant. Variant effects were quantified using the CenterMaskScorer method on the predicted DNase output. Resulting variant scores were filtered to relevant tissue ontologies to generate sequence contribution logo visualizations. To identify putative transcription factor binding sites (TFBS) disrupted or created by ISM-prioritized causal variants, we compared ISM-derived base contribution scores to TFBS motif logos in the JASPAR database^33^ accessible at: jaspar.elixir.no.

### Prioritization of linCREs for luciferase validation

Candidate elements for luciferase validation were drawn from linCREs predicted in the neural stem/progenitor cell (NSPC) Enformer context, which included 55 Hominin Gain and 23 Hominin Loss elements. We ranked candidates by a composite priority score computed as the mean of four min-max-normalized features across the candidate set: (i) the number of biosamples in which the element was called a differentially accessible linCRE, (ii) the number of brain biosamples in which it was called a linCRE, (iii) the number of predicted target genes for each element identified in the EpiMap enhancer-gene linkings^12^, and (iv) the number of target genes that are also human-ape brain differentially expressed genes^25^.This strategy aimed to prioritize elements with broad regulatory activity in the brain with functionally-relevant target genes with species-specific enhancer function in NSPCs. We designed cloning primers proceeding from the top of this ranked list, successfully cloning 12 human/chimpanzee ortholog pairs from this prioritization scheme.

We additionally included three Hominin Gain elements (HomininGain1360, HomininGain0890, HomininGain3174) predicted in a developing brain context (rather than NSPCs) that were linked to notable disease-associated genes (*CACNA1C*, *GRID1*, and *GRIN3B*), yielding 15 total ortholog pairs. None of these three developing brain elements exhibited luciferase activity in cultured NSPCs.

### iPSC culture and NSPC differentiation

Induced pluripotent stem cell (iPSC) lines were propagated feeder-free in mTeSR Plus (STEMCELL Technologies, cat #100-0276). Plates for stem cell maintenance and for neural stem/progenitor cell (NSPC) differentiation were precoated with Geltrex (Gibco, cat #A1413202). iPSC lines were differentiated into NSPCs using the StemDiff SMADi Neural Induction Kit (STEMCELL Technologies, cat #08582) following the manufacturer’s instructions. Briefly, 12 × 10^6^ iPSCs were plated in a 100 mm plate in 12 mL of Neural Induction Media (NIM) + 1 µM thiazovivin (Tocris, cat #3845) on day 0 (D0). After D0, media was changed each day to fresh NIM without thiazovivin. Cells were frozen in 1 mL of 90% NIM + 1 µM thiazovivin and 10% dimethyl sulfoxide (DMSO) at day 7 (D7) with a minimum of 2 × 10^6^ cells per cryovial. Cells were subsequently thawed for usage in luciferase assays and for immunofluorescence as detailed below. Cells were maintained in a 5% CO_2_ incubator at 37°C. Two human iPSC lines, H20961^34^ and NCRM5^35^, and three chimpanzee iPSC lines, C3651, C3649, and C886^36^, were used.

### Immunofluorescence

For immunofluorescence staining, ∼2 × 10^6^ frozen NSPCs at D7 were thawed into a precoated well of a 6-well plate in 2 mL NIM + 1 µM thiazovivin. Fresh NIM without thiazovivin was added each day. 0.6-1.2 × 10^6^ or 0.22-0.44 × 10^6^ cells were split onto coverslips on D12 in 6-well or 12-well plates, respectively. On D14, cells were washed with 1 mL of 1x phosphate-buffered saline (PBS) and fixed with 2 mL of 4% paraformaldehyde (PFA) for 10 minutes at room temperature. Coverslips were then washed for 3 × 5 minutes in 1 mL of 1x PBS, incubated in 1mL of blocking solution (8% donkey serum, 0.3% bovine serum albumin (BSA), 0.4% Triton X- 0, 0.02% sodium azide in 1x PBS) for 1 hour at room temperature, washed again for 2 × 5 minutes in 1 mL of 1x PBS, and incubated with 500 µL or 200 µL primary antibodies (mouse anti-Nestin [Sigma Aldrich, cat #MAB5326 or abcam, cat #ab22035], rabbit anti-Pax6 [Biolegend, cat #NC1586757], and goat anti-Sox2 [R&D, cat #AF2018SP]) diluted in blocking solution (1:500 [#MAB5326, #NC1586757, #AF2018SP] or 1:100 [#ab22035]) overnight at 4°C in the 6-or 12-well plates, respectively. The next day, slides were washed for 3 × 5 minutes in 1 mL of 1x PBS, and incubated with secondary antibodies (Donkey Anti-Mouse IgG H&L (Alexa Fluor® 488) [abcam, cat#ab150105], Donkey Anti-Rabbit IgG H&L (Alexa Fluor® 594) [abcam, cat #ab150076], Donkey Anti-Goat IgG H&L (Alexa Fluor® 647) [abcam, cat #ab150131]) diluted in blocking solution (1:500) for 1.5-2 hours at room temperature, and washed for 3 × 5 min in 1 mL of 1x PBS. 4′,6-diamidino-2-phenylindole (DAPI) mounting media (Invitrogen, cat #P36935) was added to the coverslip, and images were taken on an Echo Revolve microscope. All cell lines showed positive signal for the tested markers with some variability between lines (**Extended Data Fig. 9a**).

### Luciferase reporter assays

To clone plasmids for enhancer reporter assays, we amplified genomic regions of interest with polymerase chain reaction (PCR) and cloned them into the multiple restriction site in pNL3.2 (Promega, cat #N1041). We used the same primers to amplify from both human DNA (cell line: H20961) and chimpanzee DNA (cell line: C8861 or C3649). To test variant effects on enhancer activity, we mutagenized plasmids using the Mut Express II Fast Mutagenesis Kit V2 (Vazyme, cat# C214-02) per the manufacturer’s instructions. Mutagenesis primers were designed using the manufacturer’s software (https://tool.vazyme.com:18002/cetool/en-us/singlepoint.html). A complete list of cloning and mutagenesis oligonucleotides are provided in **Supplemental Table 15**.

To perform luciferase assays, NSPCs frozen at D7 were thawed into a precoated well of a 6-well plate in 2 mL NIM + 1 µM thiazovivin (D7). Fresh NIM without thiazovivin was added each day. On D12, NSPCs were split and plated at 2-4 × 10^4^ cells/well in 96-well plates in 75 µl of NIM + 1 µM thiazovivin. The next day, NSPCs were moved to 75 µL of fresh NIM and transfected with 0.2 µL Lipofectamine Stem Transfection Reagent (Thermo Fisher, cat #STEM00003), 10 ng of pGL4.53 (Promega, cat #E5011), and 190 ng of pNL3.2 or pNL3.2 containing a test genomic region in 10 µL of Opti-MEM (Thermo Fisher, cat #31985070). 24 hours later, luciferase activity was assessed with the Nano-Glo Dual Luciferase Reporter Assay System (Promega, cat #N1620) on the BioTek Synergy Neo2 (Agilent) or the SpectraMax ID5e (Molecular Devices). Enhancer reporter assays were performed in NSPCs differentiated from four different iPSC lines, C3651, C8861, H20961, and NCRM5 with three technical replicates per plasmid in each experiment.

### Luciferase data analysis

For quality control, we first filtered our luciferase measurements by calculating the mean (µ) and standard deviation (σ) of the luminescence signal across a minimum of 41 background control wells per plate (including empty wells, untransfected NSPCs, GFP-only transfected NSPCs, and pGL4.53-only transfected wells for NanoLuciferase background). We retained test wells only if they satisfied the following background threshold criteria:

1. *S_firefly_> μ_firefly +_* 6σ_firefly_
2. *S_nano_> μ_nano +_* 7σ*_nano_*
3. At least 2 of the 3 technical replicates for a given plasmid passed criteria 1 and 2.

The values 6 and 7 ensured that the signal across all test wells that passed filtering was higher than the maximum background value across all plates. Additionally, if pNL3.2 (control plasmid without a test region) did not meet the criteria above, the entire plate was excluded from the analysis.

NanoLuciferase activity (*S_nano_*) was normalized to firefly luciferase activity (*S_firefly_*) for each well to establish an activity ratio (R = *S_nano_*_/_ *S_firefly_*). For each test plasmid *i*, we calculated the mean ratio across technical replicates (*R_i_*) per plate.

To account for variable concentration and construct size, we calculated a concentration factor *C_i_* and a length factor *L_i_* as follows:

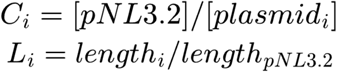

The final normalized luciferase activity *A_i_* for each test plasmid within a given experiment was calculated as the product of the mean signal normalized to pNL3.2, the concentration factor, and the length factor:

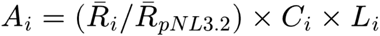

Wells were excluded if the coefficient of variation across technical replicates exceeded 0.35 or if the normalized score was non-positive. Technical replicates were collapsed to a single value per construct within each plate and species prior to statistical testing.

### Enhancer activity

For each element and species, we tested whether normalized activity exceeded background using a one-sided one-sample t-test on log2-transformed activity against a null of activity = 0 (corresponding to an activity of 1 without log transformation). An element was classified as active if it showed significant activity (BH-adjusted P < 0.05 in either species). 9 of 15 tested elements met this criterion and were retained for downstream comparisons.

### Species skew

For each active element, human and chimpanzee orthologs were compared as paired measurements within the same plate. We computed the per-plate log_2_(human/chimpanzee) activity ratio. As our deep learning predictions provided *a priori* directional hypothesis, we tested for species differences using a one-sided one-sample t-test of the paired log_2_ ratios against 0. Specifically, predicted Hominin Gain-linCRes were tested for human-specific activity (log_2_ ratio > 0), and predicted Hominin Loss-linCREs were tested for chimpanzee-specific activity (log_2_ ratio < 0). Elements with BH-adjusted P < 0.05 were considered to show significant species skew. This within-plate paired ratio test was designed to control for batch effects that confound direct comparison across plates and NSPC species of origin. For HomininGain2145 mutant constructs, we performed one-sided t-tests in the presence of clear directional hypotheses (such as Hu / Hu_T→C_ > 1) and performed a two-sided test otherwise (including Hu_T→C_ / Ch and Hu / Ch_C→T_).

---

**Supplementary Figure 1.**
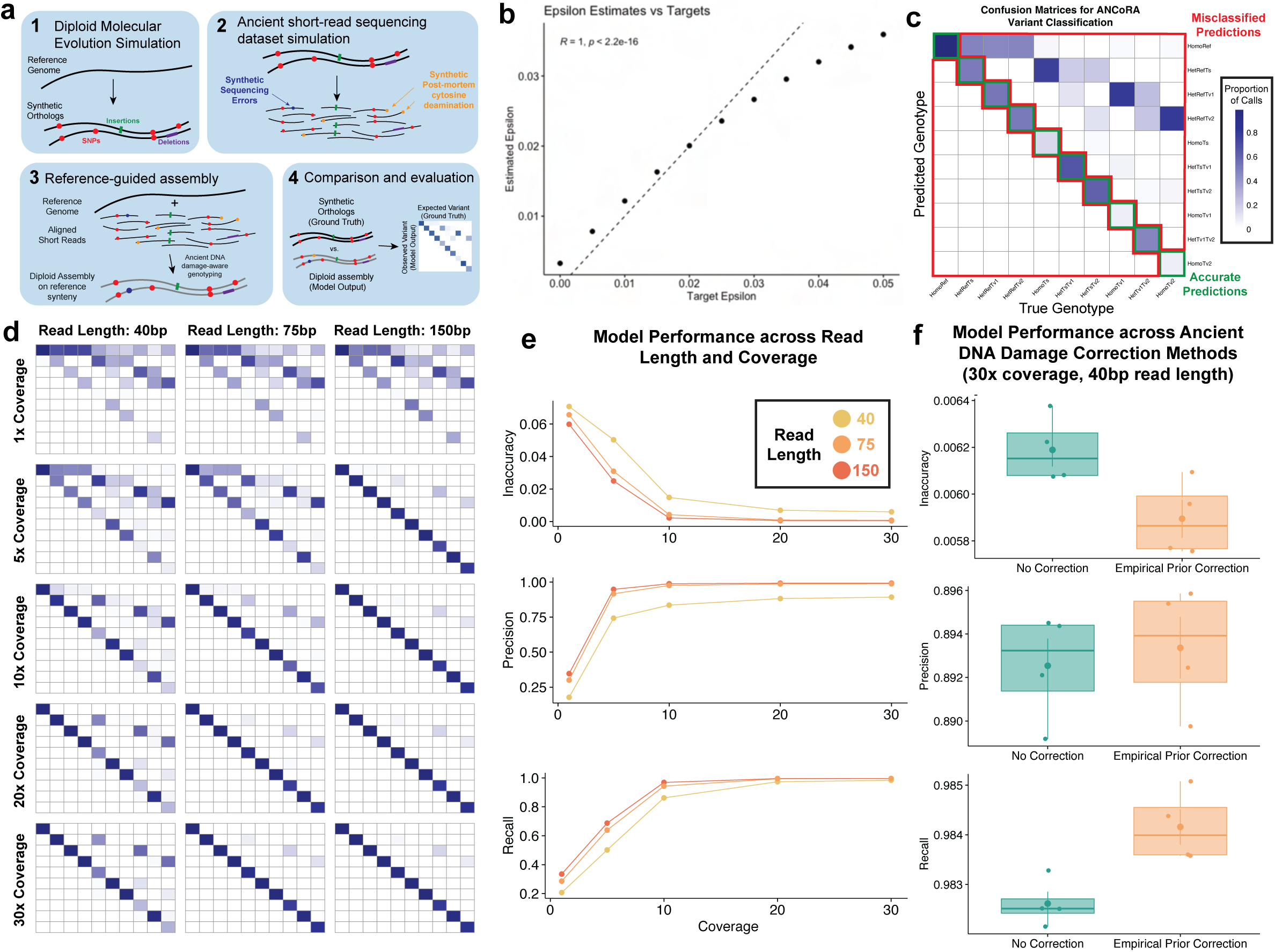

**Supplementary Figure 2.**
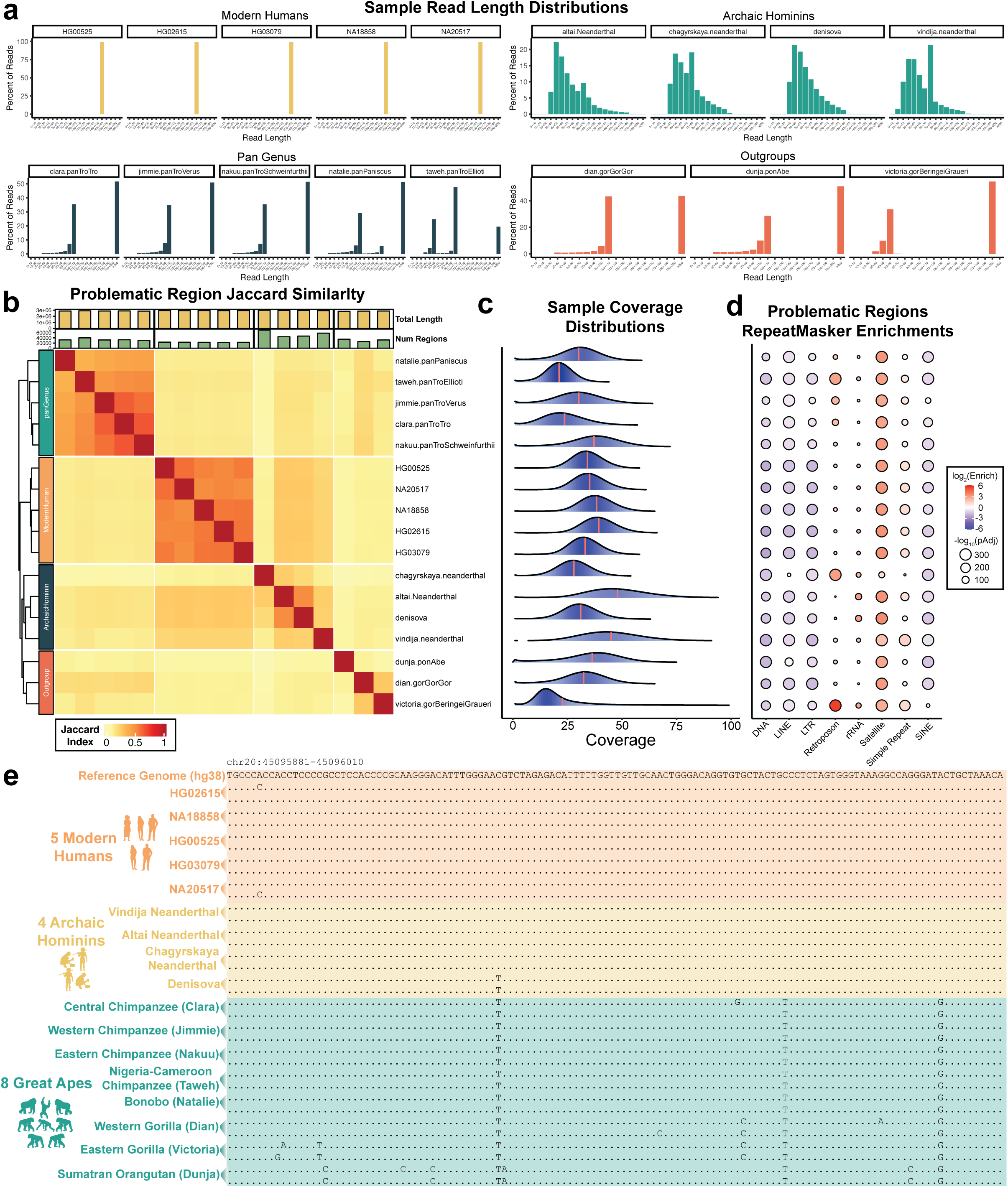

**Supplementary Figure 3.**
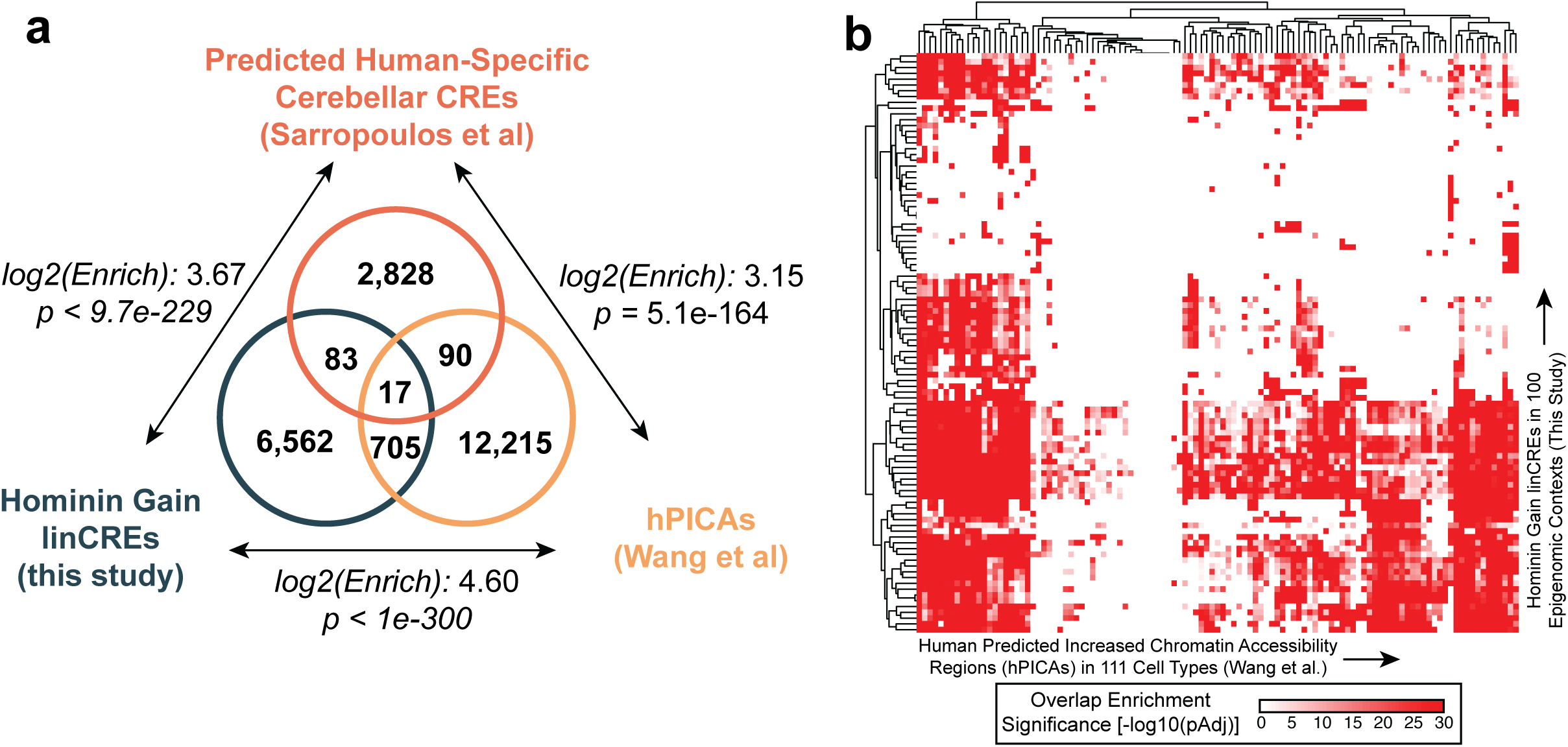

