## Supplementary Materials for "Sequence-to-function deep learning decodes human *cis*-regulatory evolution"

### Supplementary Note I: A reference-guided assembly procedure for ancient DNA short-read libraries

#### Results

To generate input sequences for sequence-to-function prediction across modern humans, archaic hominins, and great apes, we required a procedure capable of producing contiguous, personalized diploid genome assemblies from short-read sequencing libraries. Because ultra-short read lengths and postmortem damage preclude conventional overlap-consensus *de novo* assembly of high-coverage ancient DNA libraries, we adopted a reference-guided strategy[1], in which sequencing reads are aligned to a reference genome, diploid variants are called at each position, and the reference sequence is modified to reflect single-nucleotide polymorphism (SNP) and short insertion/deletion (INDEL) variants supported by the read data. The resulting assemblies retain reference synteny, which restricts our analysis to the regulatory consequences of SNPs and short INDELs and excludes structural and karyotypic variation from consideration. We view this scoping as appropriate given that current sequence-to-function models are most reliable for predicting local sequence determinants of regulatory element activity[2], and exhibit limited accuracy for long-range regulatory dependencies[3–5]. Reference-guided assembly also avoids the difficulties of *de novo* assembly from ancient libraries.

We implemented this procedure as ANCoRA (<u>A</u>ncient-DNA <u>N</u>ucleotide-damage <u>Co</u>rrection and <u>R</u>eference-guided <u>A</u>ssembly), a tool distributed as part of Gonomics, an open-source initiative for genomics software in the Go programming language[6] (**Supplementary Fig. 1a**). Rather than aiming for maximum performance in variant calling, we designed ANCoRA to favor conservative, reference-biased variant calls, since this bias minimizes overestimation of functional differences between sequences in regions of poor coverage during downstream prediction.

Ancient DNA libraries exhibit a characteristic excess of C -> T (and G -> A) misclassifications produced by postmortem cytosine deamination[7–9], which is not modeled by general-purpose variant callers such as the Genome Analysis Toolkit[8,10] or SAMtools[11] and which can bias genotyping, particularly for low-coverage samples and at heterozygous sites. Several variant callers explicitly accommodate ancient DNA damage[12,13]. ANCoRA adopts an empirical Bayes approach analogous to that of mapDamage2[14]: a per-library deamination rate is estimated directly from input pileups and incorporated into a per-site genotype likelihood (**Methods**). Applied to synthetic libraries with known damage parameters, ANCoRA effectively recovers simulated damage parameters when deamination rates were low, though increasingly underestimates deamination rates as they increase (**Supplementary Fig. 1b**).

To assess whether ANCoRA-derived assemblies are sufficiently accurate for downstream regulatory inference, we constructed a simulation-based benchmark (**Supplementary Fig. 1a**). We first generated a synthetic reference genome and introduced sequence divergence through forward evolution, mirroring the variation that distinguishes any personalized genome from a reference. We then simulated paired-end short-read libraries from these divergent haplotypes, incorporating both flat sequencing error and read-end-biased cytosine deamination characteristic of ancient DNA[9]. Synthetic reads were aligned back to the original reference, ANCoRA was applied to generate diploid assemblies, and the reconstructed haplotypes were compared base-by-base to the ground-truth divergent haplotypes (**Methods**).

Across simulations, ANCoRA recovered ground-truth genotypes with accuracy that scaled, as expected, with both sequencing coverage and read length (**Supplementary Fig. 1c-e**). At the expected library conditions for ancient DNA (30x coverage, 40bp average read length, ancient DNA damage parameter = 0.05, and a flat sequencing error rate of 0.01), we achieved a total substitution accuracy of 99.4%. Including the empirical damage prior modestly improved overall accuracy at short read lengths (99.37 -> 99.4%), primarily by increasing recall (**Supplementary Fig. 1f**). These results gave us confidence that ANCoRA-derived assemblies are reliable inputs for sequence-to-function prediction, and motivated our subsequent restriction to high-coverage archaic libraries (**Supplementary Note 2**).

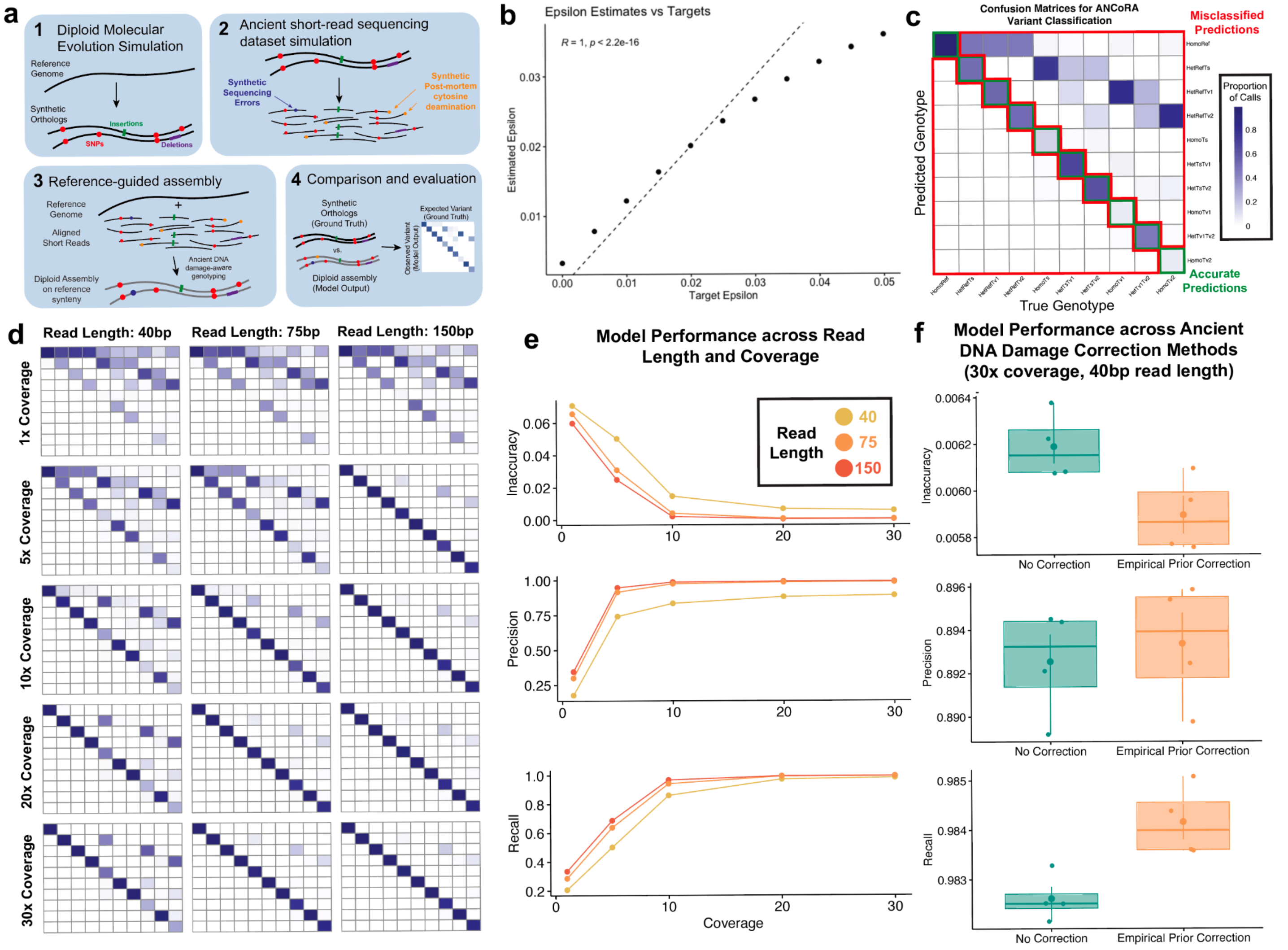

### Supplementary Figure 1. ANCoRA model validation.

(**a**) Workflow for ANCoRA validation, including forward evolution simulation, ancient DNA short-read simulation, reference-guided assembly, and evaluation.

(**b**) Evaluation of estimation of ancient DNA cytosine deamination rates from synthetic short read libraries. Target rates correlate strongly with ANCoRA-estimated rates.

(**c**) Example confusion matrix for ANCoRA variant classification. Diploid genotypes are organized into 10 categories, conditional on the reference base: HomoRef (homozygous reference), HetRefTs (heterozygous reference/transition), HetRefTv1 (heterozygous reference/transversion 1, referring to the first transversion, alphabetically; ex. Ref=A, Geno=AC), HetRefTv2 (heterozygous ref transversion 2, ex.Ref=A, Geno=AT), HomoTs (homozygous transition), HetTsTv1 (heterozygous transition/transversion 1), HetTsTv2 (heterozygous transition/ transversion 2), HomoTv1 (homozygous transversion 1), HetTv1Tv2 (heterozygous transversion 1/transversion 2), and HomoTv2 (homozygous transversion 2). Matrix entries indicate the proportion of bases for each true genotype called as each predicted genotype. Green boxes indicate accurate predictions, while red boxes highlight misclassified positions.

(**d**) Confusion matrices for ANCoRA performance across read lengths (40, 75, 150bp) and sequencing coverage (1x, 5x, 10x, 20x, 30x).

(**e-f**) ANCoRA model performance, evaluated by inaccuracy (top), precision (middle), and recall (bottom).

(**e**) ANCoRA performance metrics as a function of sequencing coverage, colored by read length. Model performance improves with both read length and coverage.

(**f**) Metrics displayed with and without an empirical prior correction for ancient DNA damage at 30x coverage and 40bp read length. Correction slightly improves inaccuracy by increasing recall.

### Methods

#### ANCoRA overview

The Ancient-DNA Nucleotide-damage Correction and Reference-guided Assembly (ANCoRA) tool generates reference-guided unphased diploid assemblies from aligned short reads. ANCoRA takes as input a reference genome in FASTA format and a sorted BAM file of aligned reads from single- or paired-end libraries, generates per-site pileups, and infers diploid genotypes (substitutions and short INDELs) within an empirical Bayes framework. Output assemblies are produced by introducing called variants into the reference sequence while maintaining reference synteny; heterozygous variants are assigned to the two output haplotypes at random, and the resulting assemblies therefore represent unphased haplotypes.

This tool relies on several assumptions chosen to keep inference tractable for our application. Input reads are assumed to be derived from diploid cells with no copy-number variation. Sites are treated as independent under a time-reversible single-site mutation model, and sequencing errors are modeled with uniform classification rates across bases (Jukes-Cantor). The reference genome serves as an informative prior: in the absence of supporting reads, ANCoRA emits the reference allele at each position. Combined with the conservative damage model described below, this yields reference-biased output, which we view as desirable for downstream sequence-to-function prediction since it reduces the rate at which low-confidence calls propagate into spurious functional differences. ANCoRA is implemented in Go with minimal external dependencies as part of the Gonomics project[6].

#### Genotype Likelihoods

**Substitutions.** ANCoRA calls genotypes from pileups by computing posterior probabilities following standard formulations[10–12]:

$$P(Geno|Pile) = \frac{P(Pile|Geno)P(Geno)}{P(Pile)}$$

For diploid sites, $Geno \in \{AA, AC, AG, AT, CC, CG, CT, GG, GT, TT\}$. For haploid sites, $Geno \in \{A, C, G, T\}$. Observed base counts at a position are denoted $\{a, c, g, t\}$.

For an example pileup such as $[a:1, c:6, g:17, t:0]$, evidence for three distinct bases at a single diploid site is accommodated by an error term $\epsilon$, representing the cumulative effects of PCR amplification, sequencing error, and alignment artifact. The likelihood of an observed pileup under a given genotype is modeled as a multinomial over the four bases $(k = 4)$. For the homozygous genotype $AA$:

$$p(Pile|AA) = \frac{n!}{a!c!g!t!}(1-\epsilon)^a(\frac{\epsilon}{3})^{c+g+t}$$

where $\epsilon/3$ reflects the equal probability of misclassification to each of the three incorrect bases. For a heterozygous genotype, e.g. $AC$:

$$p(Pile|AC) = \frac{n!}{a!c!g!t!}(0.5 - \frac{\epsilon}{3})^{a+c}(\frac{\epsilon}{3})^{g+t}$$

The coefficient $0.5 - \epsilon/3$ can be derived by considering both correct base observations and misclassifications. For a heterozygous AC genotype, the probability of correctly observing A from the A haplotype is $0.5 \times (1 - \epsilon)$, assuming both haplotypes are sampled with equal probability. Additionally, there is a chance for a base sampled from the C haplotype to be misclassified as A, contributing to the observed count of A with frequency $\epsilon/3$. Thus, the total probability of observing A in the pileup is:

$$0.5 \times (1 - \epsilon) + 0.5 \times \frac{\epsilon}{3}$$

Because all downstream comparisons reduce to likelihood ratios, the multinomial coefficient is never explicitly evaluated.

**Extension for ancient DNA damage.** Postmortem hydrolytic decomposition produces characteristic C to T misclassifications (and G to A misclassifications on the reverse strand of double-stranded libraries)[15]. We extend the per-site likelihood with a second error term, $\lambda$, representing the per-base deamination rate, while $\epsilon$ retains its earlier role as a flat baseline misclassification error rate. Under this two-parameter model, the conditional probability of observing each base in a read given the underlying base in the source DNA is:

| | | **Base in Organism** | | | |
|---|---|---|---|---|---|
| | | **A** | **C** | **G** | **T** |
| **Base in Sequencing Read** | **A** | $1 - \epsilon$ | $\frac{\epsilon}{3}$ | $\frac{\epsilon}{3} + \lambda$ | $\frac{\epsilon}{3}$ |
| | **C** | $\frac{\epsilon}{3}$ | $1 - \epsilon - \lambda$ | $\frac{\epsilon}{3}$ | $\frac{\epsilon}{3}$ |
| | **G** | $\frac{\epsilon}{3}$ | $\frac{\epsilon}{3}$ | $1 - \epsilon - \lambda$ | $\frac{\epsilon}{3}$ |
| | **T** | $\frac{\epsilon}{3}$ | $\frac{\epsilon}{3} + \lambda$ | $\frac{\epsilon}{3}$ | $1 - \epsilon$ |

This parameterization yields the following homozygous likelihoods:

$$p(Pile|AA) = (\frac{\epsilon}{3})^{c+g+t}(1 - \epsilon)^{a}$$

$$p(Pile|CC) = (\frac{\epsilon}{3})^{a+g}(1 - \epsilon - \lambda)^{c}(\frac{\epsilon}{3} + \lambda)^{t}$$

$$p(Pile|GG) = (\frac{\epsilon}{3} + \lambda)^{a}(\frac{\epsilon}{3})^{c+t}(1 - \epsilon - \lambda)^{g}$$

$$p(Pile|TT) = (\frac{\epsilon}{3})^{a+c+g}(1 - \epsilon)^{t}$$

For heterozygous genotypes, we assume that each chromosome is sampled with equal probability. Therefore, the probability of observing a particular nucleotide in a pileup, such as observing an A from an AC genotype, is the weighted sum of two events: 0.5 times the probability of observing an A from a chromosome containing an A, and 0.5 times the probability of observing an A from the chromosome with a C genotype:

$$P(a|AC) = 0.5 \cdot P(a|A) + 0.5 \cdot P(a|C)$$

Thus, the diploid heterozygous genotype likelihood functions under this parameterization are expressed as follows:

$$p(Pile|AC) = (0.5 - \frac{\epsilon}{3})^a (0.5 - \frac{\epsilon}{3} - \frac{\lambda}{2})^c (\frac{\epsilon}{3})^g (\frac{\epsilon}{3} + \frac{\lambda}{2})^t$$

$$p(Pile|AG) = (0.5 - \frac{\epsilon}{3} + \frac{\lambda}{2})^a (\frac{\epsilon}{3})^{c+t} (0.5 - \frac{\epsilon}{3} - \frac{\lambda}{2})^g$$

$$P(Pile|AT) = (0.5 - \frac{\epsilon}{3})^{a+t} (\frac{\epsilon}{3})^{c+g}$$

$$p(Pile|CG) = (\frac{\epsilon}{3} + \frac{\lambda}{2})^a (0.5 - \frac{\epsilon}{3} - \frac{\lambda}{2})^{c+g} (\frac{\epsilon}{3} + \frac{\lambda}{2})^t$$

$$p(Pile|CT) = (\frac{\epsilon}{3})^{a+g} (0.5 - \frac{\epsilon}{3} - \frac{\lambda}{2})^c (0.5 - \frac{\epsilon}{3} + \frac{\lambda}{2})^t$$

$$p(Pile|GT) = (\frac{\epsilon}{3} + \frac{\lambda}{2})^a (\frac{\epsilon}{3})^c (0.5 - \frac{\epsilon}{3} - \frac{\lambda}{2})^g (0.5 - \frac{\epsilon}{3})^t$$

**Insertions.** Pileup depth is computed as $n = a + c + g + t$. Observed insertions are stored in a hash map keyed by insertion sequence, with values giving observed counts. Multiple distinct insertions can be observed at the same site (e.g. complex heterozygous insertions or sequencing errors). To keep the model tractable, we consider only the two most frequently observed insertions, $I_a$ and $I_b$, as candidate alleles, treating all other observed insertions as errors.

At each position, we consider the following possible genotypes: $I_aI_a$, $I_aI_b$, $I_aBase$, and $BaseBase$, where $Base$ denotes the absence of an insertion. Genotypes $I_bBase$ and $I_bI_b$ are excluded as exceedingly unlikely given the count ordering.

Let $i_{tot}$ denote the total count for all observed insertions, including those from less frequently observed insertions ($i_c$, $i_d$, etc.), which would not be considered in candidate genotypes. As we choose to ignore less frequently observed insertions ($i_c$, $i_d$, etc.) for genotype likelihood calculations, we exclude them from the sequencing depth estimate $n$. Therefore, we calculate $base$, the number of reads for 'no insertion', as $n - i_{tot}$.

In this program, we model the likelihoods of insertion states as a multinomial distribution with three outcomes: $I_a$, $I_b$, and $Base$. A read sampled from a chromosome with genotype $I_a$ can be misclassified with probability $\epsilon$ as either $I_b$ or $Base$ with equal probabilities ($\epsilon/2$). From here, we can derive the likelihood expressions for the following four genotypes:

$$P(Pile|I_aI_a) = (1 - \epsilon)^{i_a} (\frac{\epsilon}{2})^{base+i_b}$$

$$P(Pile|BaseBase) = (1 - \epsilon)^{base} (\frac{\epsilon}{2})^{i_a+i_b}$$

$$P(Pile|I_aI_b) = (0.5 - \frac{\epsilon}{4})^{i_a+i_b} (\frac{\epsilon}{2})^{base}$$

$$P(Pile|I_aBase) = (0.5 - \frac{\epsilon}{4})^{i_a+base} (\frac{\epsilon}{2})^{i_b}$$

Note that in our implementation, a single error rate $\epsilon$ is shared between substitutions and INDELs.

**Deletions.** We store deletion counts in a hash map with integer keys representing deletion lengths, and values corresponding to the count corresponding to that deletion length. Deletions are treated similarly to insertions, with only the two most occurring deletions ($D_a$ and $D_b$) considered in genotypes. Let $d_{tot}$ represent the total number of deletion reads, and $base$ represent the number of reads without a deletion, where $base = n - d_{tot}$.

$$P(Pile|D_aD_a) = (1 - \epsilon)^{d_a} (\frac{\epsilon}{2})^{base+d_b}$$

$$P(Pile|BaseBase) = (1 - \epsilon)^{base} (\frac{\epsilon}{2})^{d_a+d_b}$$

$$P(Pile|D_aD_b) = (0.5 - \frac{\epsilon}{4})^{d_a+d_b}(\frac{\epsilon}{2})^{base}$$

$$P(Pile|D_aBase) = (0.5 - \frac{\epsilon}{4})^{d_a+base}(\frac{\epsilon}{2})^{d_b}$$

**Haploid Base Calls in Heterozygous Deletions.** A corner case now arises in the genotype $D_aBase$, which we have illustrated with the sample alignment below:

```
Ref:  ATGGCACTATTG
HapA: ATGGC----TTG
HapB: ATGGCACGATTG
```

In this example, the first haplotype (HapA) has a four base pair heterozygous deletion relative to the reference. However, the second haplotype (HapB) has a T to G substitution in the third position of the deleted region.

To handle this case, ANCoRA makes haploid variant calls for each base in the deleted region. The likelihood function used to evaluate haploid substitutions is identical to the function used to evaluate homozygous diploid genotypes: $p(Pile|AA) = p(Pile|A)$. Similarly, haploid insertions and deletions follow the homozygous INDEL likelihood functions described earlier.

An additional case arises from complex heterozygous deletions. Consider an example where the first haplotype has a deletion of 3 base pairs and the second haplotype has a deletion of 2 base pairs. In this case, we would make a haploid call for only the third base of the second haplotype.

### Prior Probabilities

The prior $p(Geno)$ is estimated under an empirical Bayes scheme prior to variant calling. In the ANCoRA implementation, this is performed by the subcommand *ancora prior* which is run prior to the assembler subprogram *ancora build*.

**Substitution prior.** The substitution prior is represented by conditional Dirichlet distributions in a 10x4 matrix, reflecting the probability of observing each of the 10 diploid genotypes conditional on the reference base. For each position, ANCoRA initially infers the diploid genotype using a flat prior - a uniform 10x4 matrix with all values set to 0.1, reflecting equal probabilities for all genotypes. Next, a 10x4 matrix is populated with counts reflecting the empirical frequencies for each genotype given each reference base. These counts are then normalized to generate the final prior distribution.

These priors are written to human-readable text files, allowing users to edit and specify custom priors. Users may use either the pre-built flat prior or define a non-empirical prior specified by the hyperparameters $\delta$, the total divergence rate, and $\gamma$, the ratio of transitions to transversions. All experiments described in this work, unless otherwise specified, were performed with ANCoRA empirical priors.

**Error and damage rate estimation.** ANCoRA also generates maximum likelihood estimates for the error rate $\epsilon$ and the ancient error rate $\lambda$.

Consider a position with the homozygous genotype $AA$ or $TT$. Since adenine and thymine are unaffected by cytosine deamination, all observed errors at these sites can be attributed to the sequencing error rate $\epsilon$. At positions with homozygous genotypes $CC$ and $GG$, observed errors result from the joint effects of both $\epsilon$ and the deamination rate $\lambda$.

From this, we derive maximum likelihood estimates for both $\hat{\epsilon}$ and $\hat{\epsilon} + \hat{\lambda}$. The maximum likelihood estimate for the deamination rate $\hat{\lambda}$ is then calculated by subtracting $\hat{\epsilon}$ from the joint estimate: $\hat{\lambda} = (\hat{\epsilon} + \hat{\lambda}) - \hat{\epsilon}$.

$$\hat{\epsilon} = \frac{Errors in AA and TT Genotypes}{Total Bases in AA and TT Genotypes}$$

$$\hat{\epsilon} + \hat{\lambda} = \frac{Possible deaminations (C \rightarrow T in CC and G \rightarrow A in GG)}{Total Bases in CC and GG Genotypes}$$

**INDEL prior.** We treat insertions and deletions as both occurring at equal frequency based on user-specified hyperparameters $\kappa$ and $\delta$, where $\delta$ is the probability of a divergent site, and $\kappa$ is the probability that a divergent site is an INDEL. INDEL genotype prior probabilities are as follows:

$$P(I_aI_a) = P(I_bI_b) = P(D_aD_a) = P(D_bD_b) = (\kappa\delta)^2$$

$$P(I_aI_b) = 2(\kappa\delta)^2$$

$$P(I_aBase) = P(D_aBase) = 2\kappa\delta$$

$$P(BaseBase) = 1 - 4\kappa\delta - 3(\kappa\delta)^2$$

**Haploid Priors.** When making haploid INDEL calls, we use the following prior values:

$$P(I_a) = P(D_a) = \kappa\delta$$

$$P(Base) = 1 - \kappa\delta$$

**Bayesian Framework for Diploid Variant Calling From Pileups.** From Bayes' rule, we establish the probability of a genotype given a pileup to follow equation:

$$P(Genotype|Pileup) = \frac{P(Pileup|Genotype)P(Genotype)}{P(Pileup)}$$

The ratio of posterior probabilities between two genotypes, $G_a$ and $G_b$, for the same pileup can be expressed as:

$$\frac{P(G_a|Pileup)}{P(G_b|Pileup)} = \frac{P(Pileup|G_a)P(G_a)}{P(Pileup|G_b)P(G_b)}$$

For each pile, we consider 10 substitution genotype hypotheses, 4 insertion hypotheses, and 4 deletion hypotheses, for a total of 18 hypotheses.

### Simulation-based evaluation

**Forward evolution simulation.** We began by generating a synthetic reference genome, containing two chromosomes, each at a length of one megabase, using the Gonomics program *randSeq*. We next implemented the Gonomics program *simulateEvol withIndels*, which simulates forward evolution, to introduce variants to the resulting synthetic reference genome. The *simulateEvol* program introduces both single nucleotide polymorphisms (SNPs) and small insertions or deletions (INDELs) following user-specified evolutionary parameters: *-propIndel 0.1 -branchLength 0.05 -transitionBias 3 -lambda 1 -gcContent 0.42*. Here *branchLength* reflects the total degree of sequence divergence, introducing one mutation for approximately every 20 bases. We set *propIndel* to 0.1, indicating that 10% of these mutations were INDELs, with the remaining 90% of mutations as SNPs. The transition bias of 3 reflects the elevated rates of transitions (A↔G, C↔T) compared to transversions observed in human germline evolution[16]. Lambda reflects the rate parameter of an exponential distribution used to sample INDEL lengths, ensuring that most INDELs are short (e.g., single base pair), while still allowing for longer variants. To generate random genomic sequences for insertions, we specify a GC content of 0.42 to match the average GC content of the human genome.

**Short read library simulation.** We expanded the existing *simulateSam* program from the Gonomics library to generate synthetic short read sequencing datasets. This program generates synthetic Illumina-style paired-end short reads from a reference genome in FASTA format, based on user-specified parameters for read length and coverage. We performed simulations at a series of read lengths (40, 75, 150bp) and coverage levels (1x, 5x, 10x, 20x, 30x) to evaluate ANCoRA performance across diverse sequencing conditions.

In addition to the existing functionality, we added several new parameters to model sequencing errors in short reads, which can arise from both PCR amplification and the sequencing process. The *flatErrorRate* parameter was set to 0.01, such that 1% of bases in synthetic reads were misclassified. These sequencing errors were modeled as “flat” errors, following a uniform error spectrum analogous to the Jukes-Cantor mutation model.

To optimize error generation, we treated each position independently and assumed that the number of errors per read follows a binomial distribution with parameters $n = readLength$ and $p = flatErrorRate$. Using Walker’s alias method[17], we generated a binomially distributed random variate $X$ to represent the number of sequencing errors in each read. We then sampled $X$ positions without replacement from the read and introduced sequencing errors at these positions, ensuring efficient and biologically plausible error modeling.

**Simulating cytosine deamination in ancient DNA samples.** We also expanded the *simulateSam* program to model patterns of postmortem cytosine deamination characteristic of ancient DNA libraries. Cytosine deamination events are concentrated near the ends of reads, and their distance to read ends is well modeled by a geometric distribution[15]. In our model, we first defined the parameter *ancientErrorRate* to 0.05, reflecting the rate of cytosine deaminations, observed as C to T mutations or G to A mutations on the reverse strand. Thus, a read derived from a genomic position with a C in the original genome will show a T in the read at a probability based on the additive effects of $\epsilon$, the flat error rate, and $\lambda$, an ancient DNA-specific cytosine deamination rate:

$$P(T_{read}|C_{base}) = P(A_{read}|G_{base}) = \epsilon + \lambda$$

**ANCoRA performance evaluation.** We next aligned synthetic short read libraries to the initial reference genome using *bwa mem*[18] with default parameters and used ANCoRA to call variants and construct reference-guided assemblies. We used the *multiFaDir* parameter of *ancora build* to represent the resulting assemblies in *multiFa* alignment format, and used the Gonomics utility *mergeMultiFa* to construct a five-way multiple alignment comprised of the original reference genome, the two synthetic haplotypes generated by *simulateEvol*, and the two reconstructed haplotypes generated by ANCoRA. We then evaluated ANCoRA accuracy using the *ancora score* subcommand in *baseMatrix* mode. This tool compares diploid genotypes from the reconstructed haplotypes to ground truth synthetic haplotypes on a per-base level. Treating diploid genotypes as unphased, each position in the ground truth synthetic sequence and the ANCoRA reconstruction can be described as one of ten genotypes: $Geno \in \{AA, AC, AG, AT, CC, CG, CT, GG, GT, TT\}$. Thus, *ancora score* constructs a 10x10 confusion matrix, and total inaccuracy as the proportion of off-diagonal elements. We calculated precision as the proportion of true positive predictions among all positions predicted as variants, excluding positions with a ground truth genotype of homozygous reference. Similarly, recall was defined as the proportion of variant sites (positions with a ground truth genotype other than homozygous reference) that were correctly classified by ANCoRA.

### Supplementary Note II: Generation of personalized assemblies and a 35-way hominin and great ape alignment

#### Results

To survey *cis*-regulatory sequence divergence across hominin and great ape evolution, we used ANCoRA to construct personalized reference-guided assemblies from short-read sequencing libraries. Based on our findings that ANCoRA performance was highly dependent on sequencing coverage, we accessed the four available archaic hominin samples that have been sequenced to high coverage, including one Denisovan[19] and three Neanderthals[20–22]. Additionally, we used ANCoRA to generate diploid reference-guided assemblies from five modern humans[23] as well as 4 chimpanzees, 1 bonobo, 2 gorillas, and 1 orangutan from the Great Ape Genome Project[24]. As expected, archaic hominin sequencing libraries exhibit shorter read lengths, reflecting the barriers against *de novo* assembly of ancient DNA (**Supplementary Fig. 2a**). All included libraries were sequenced to high coverage (**Supplementary Fig. 2c**).

Because all four high-coverage archaic samples are derived from putatively female individuals, we restricted modern human and great ape selection to female samples to maintain consistent diploid coverage of the X chromosome and to permit exclusion of the Y chromosome from all downstream analyses. The single exception is the *Pan troglodytes ellioti* specimen Taweh, previously reported to exhibit an XXY karyotype[24]. Modern human libraries were drawn from the 1000 Genomes Project[23] to span global genetic diversity.

To mitigate potential genotyping biases in repetitive regions and other regions of poor mappability, we defined problematic regions for each assembly as regions with unusually high read coverage (**Supplementary Fig. 2b**). We encountered similar problematic regions across samples, which were highly enriched for satellites, simple repeats, and rRNA gene arrays (**Supplementary Fig. 2d**). Problematic regions for each library were masked to the reference sequence in the final assemblies.

Because each ANCoRA assembly carries its own short INDELs relative to the reference, each assembly has its own coordinate system. We therefore aggregated the 34 reference-guided assemblies, together with the hg38 reference, into a 35-way genome-wide multiple alignment maintained on shared hg38 coordinates (**Supplementary Fig. 2e**). This alignment forms the input for genome-wide sequence-to-function prediction described in the main text.

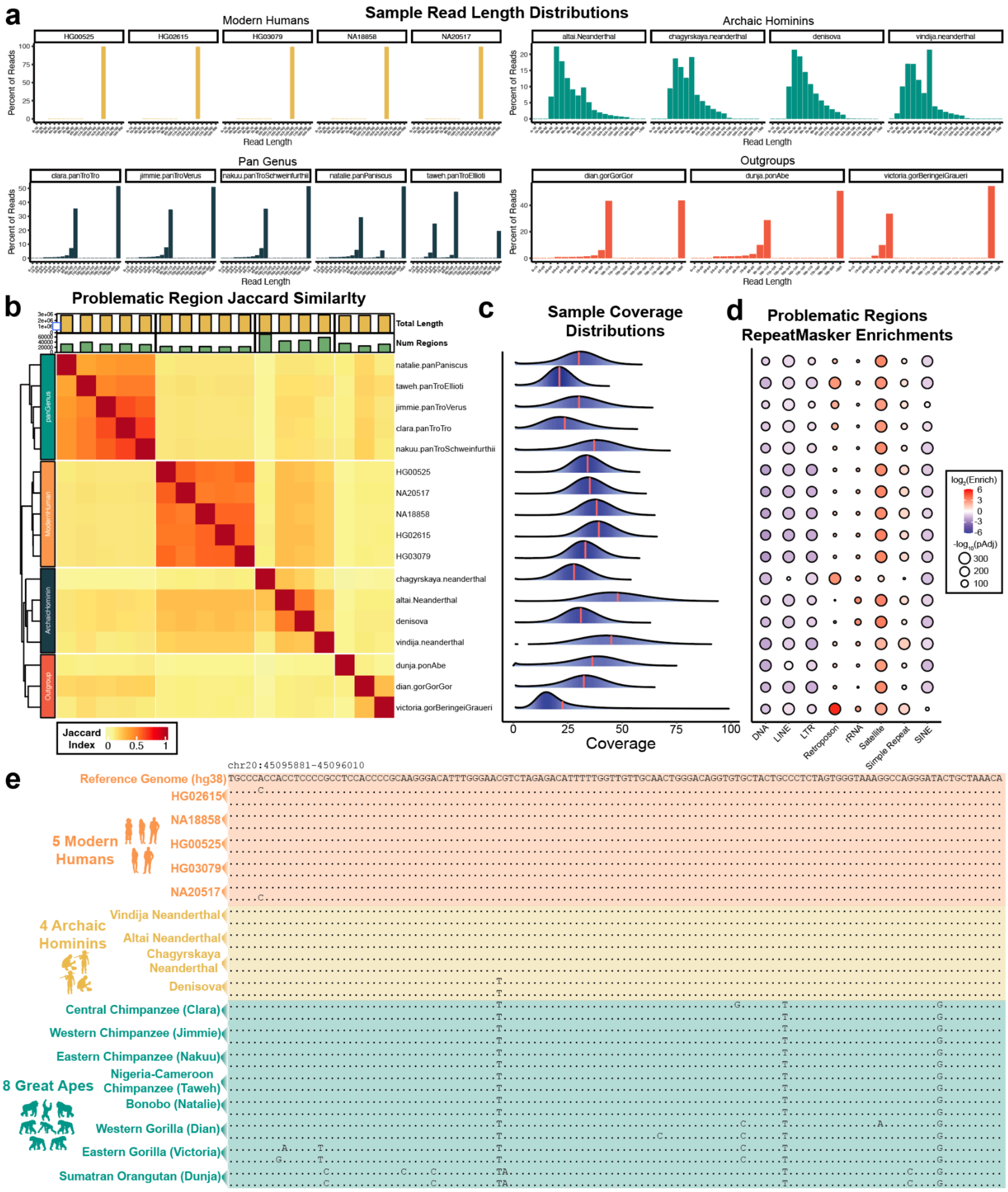

**Supplementary Figure 2. Library composition and characterization of problematic regions in reference-guided assemblies.**

(**a**) Read length distributions across samples. Archaic hominin samples exhibit significantly shorter average read lengths and greater read length variability.

(**b**) Jaccard similarity matrix between the problematic regions in each library. Column annotations display the total number of bases in problematic regions (Total Length) and the number of distinct problematic regions (Num. Regions).

(**c**) Coverage distributions for each sequencing library. Red line marks the mean coverage for each library.
(**d**) Overlap enrichments between problematic regions of each sequence and *repeatMasker* repetitive element annotations. Problematic regions are enriched for satellite, simple repeats, and rRNA gene repeats across samples.

(**e**) Segment of a genome-wide 35-way multiple alignment of ANCoRA-derived assemblies.

### Methods

#### Generation of modern human, archaic hominin, and great ape diploid assemblies with ANCoRA

We used ANCoRA to generate reference-guided assemblies using short read sequencing libraries from five modern humans, four archaic hominins, and eight great apes. Only four archaic hominin samples have been sequenced to high coverage, as most ancient DNA samples are of insufficient quality for such sequencing. All four are included in this analysis: one Denisovan[19] and three Neanderthals - Altai[20], Vindija[21], and Chagyrskaya[22]. Five modern humans, representing diverse global populations, were selected from the 1000 Genomes Project, including: HG02615 (Gambian in Western Division, The Gambia - Mandinka), NA18858 (Yoruba in Ibadan, Nigeria), HG00525 (Han Chinese South), HG03079 (Mende in Sierra Leone), and NA20517 (Toscani in Italy)[23]. Great ape libraries were accessed from the Great Ape Genome Project[24] and included the chimpanzee specimens Clara (*Pan troglodytes troglodytes*), Jimmie (*Pan troglodytes verus*), Nakuu (*Pan troglodytes schweinfurthii*), Taweh (*Pan troglodytes ellioti*), the bonobo specimen Natalie (*Pan paniscus*), the gorilla specimens Dian (*Gorilla gorilla gorilla*) and Victoria (*Gorilla beringei graueri*), and the orangutan specimen Dunja (*Pongo abelii*).

Incidentally, all four archaic hominin samples are derived from putative biological females. To maintain consistency, we restricted our analysis of modern humans and great apes to female samples, allowing for diploid coverage of the X chromosome, excluding the Y chromosome from all analyses. One exception in our analysis was the Pan troglodytes ellioti specimen 'Taweh', which was previously reported to exhibit an XXY karyotype[24].

All sequencing libraries were accessed in either BAM or CRAM format. CRAM format files were first converted to BAM format (*samtools view*)[11] and reverted to FASTQ-format files (*bedtools bamtofastq*)[25]. We then aligned reads to a version of the human reference genome GRCh38 (hg38) modified to include only canonical chromosomes and excluding alternative haplotypes, unplaced contigs, and the mitochondrial chromosome using *bwa-mem*[18,26].

**Read length and coverage distributions.** We implemented *samInfo* in the Gonomics library to assess the genome-wide distribution of sequencing coverage from an input sequencing library. This program takes as input a short read sequencing dataset in SAM/BAM alignment format and generates a sequencing coverage histogram from pileups. We also implemented the *samInfo readLength* subcommand to calculate read length distribution histograms from input SAM/BAM files.

**Analysis of problematic regions.** To mitigate potential genotyping biases in repetitive regions with poor mappability, we defined problematic regions for each sample as regions with unusually high coverage identified from empirical coverage distributions. To this end, we developed the program *samCoverage*, available as part of the Gonomics repository[6]. This program calculates empirical coverage distributions from SAM/BAM format short read datasets, fits the resulting distribution to a Poisson distribution, and defines a threshold coverage depth at a user-specified proportion of the cumulative distribution function. In this analysis, we defined the set of genomic regions with the highest 0.1% of coverage as problematic. We then used the Gonomics program *wigTools peaks* to identify these regions in each sample.

To determine if similar genomic regions were identified as problematic regions across genome assemblies, we analyzed the Jaccard Index between sets of problematic regions from each assembly using *bedtools jaccard*[25]. We visualized the resulting Jaccard matrix with the pheatmap R package (v1.0.12) (**Supplementary Fig. 2b**). Next, we determined overlap enrichments between problematic regions and repetitive genomic elements using the *gonomics overlapEnrichments* tool[6] with repeatMasker element annotations accessed from the UCSC Genome Browser[27] (**Supplementary Fig. 2d**).

We then used ANCoRA to generate reference-guided assemblies from these libraries using an ANCoRA-derived empirical prior matched to each sample. Problematic regions were masked to the hg38 reference sequence for each assembly using the Gonomics utility *multiFaSequenceSwap*.

**Genome-wide multiple alignment.** We used the *multiFaDir* option of ANCoRA to generate reference-guided assemblies in *multiFa* alignment format, which results in three-way alignments of the *hg38* reference genome together with the diploid reference-guided assemblies. We used the Gonomics tool *mergeMultiFa*, which merges multiFa format alignments on a shared reference genome, to construct a genome-wide 35-way alignment including the *hg38* human reference genome and 34 haplotypes across 17 individuals spanning modern humans, archaic hominins, and great apes. We used the Gonomics program *multFaVisualizer* to visualize distinct genomic regions across this alignment, as in **Supplementary Fig. 2e**.

**Coordinate interoperability.** As we model short INDELs in addition to SNPs, each ANCoRA assembly has distinct genome coordinates. To interoperate across assemblies, we first used *faFilter* from the Gonomics library to subset multiFa format alignments into pairwise alignments. Then, we used *multiFaToChain* to generate chain format files, facilitating conversion of BED format region files across assembly coordinates with *liftOver*[27]. We used the UCSC tool *chainSwap* to produce additional chain files when mapping regions on assembly-specific coordinates to *hg38* reference coordinates.

### Supplementary Note III: Comparison with concurrent studies leveraging sequence-to-function models for evolutionary prediction

As this study neared submission, two concurrent studies were published applying sequence-to-function deep learning methods to predict lineage-specific regulatory elements in human evolution. Sarropoulos et al. a model to predict human-specific *cis*-regulatory elements in the developing cerebellum[28]. Wang et al. trained 111 independent, cell-type-specific convolutional neural networks on single-cell ATAC-seq data to identify “human predicted increased chromatin accessibility regions” (hPICAs) by comparing the modern human reference genome to an inferred human-chimpanzee ancestor[29].

Despite differences in model architectures, training datasets, input sequences, and statistical frameworks, we observed highly significant overlap enrichments across all prediction sets (**Supplementary Fig. 3a**). Our predicted Hominin Gain-linCREs were enriched for both the cerebellar CREs predicted by Sarropoulos et al. ($\log_2$(Enrich) = 3.67, $p < 9.7 \times 10^{-229}$) and the hPICAs predicted by Wang et al. ($\log_2$(Enrich) = 4.60, $p < 1 \times 10^{-300}$). While these enrichments are highly significant, the absolute number of overlapping elements constitutes a minority of the distinct loci in each dataset: 17 elements were identified as human-gained in all three studies, and 6,562 Hominin Gain-linCREs were not previously identified by Wang or Sarropoulos (89.1%).

We hypothesized that the difference in prediction sets is partially driven by differences in the sampled epigenomic contexts. Supporting this, overlap enrichments between Hominin Gain-linCREs, stratified by the 100 Enformer biosample prediction contexts and hPICAs, stratified by the 111 cell types analyzed by Wang et al. indicates that concordance between our prediction sets is driven by only a subset of epigenomic contexts (**Supplementary Fig. 3b**).

In contrast to these studies, the present work was designed to consider a broader range of *cis*-regulatory turnover, encompassing human-specific loss of ancestral regulatory function, as well as regulatory differences between modern humans and archaic hominin. Rather than comparing modern human and great ape reference genomes or ancestral sequences inferred from reference genome, our approach predicts accessibility across personalized genomes, enabling us to disentangle intraspecific polymorphism from fixed species differences, which have historically confounded comparative studies[30].

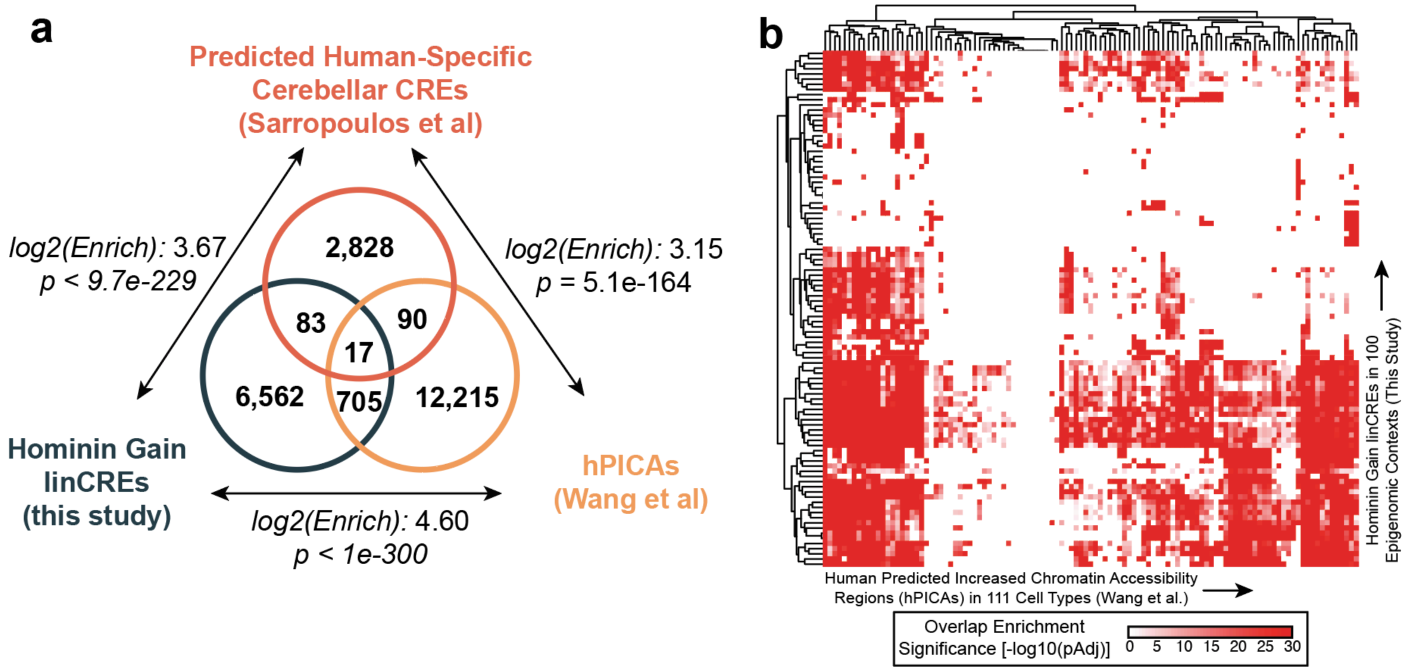

**Supplemental Figure 3: Comparison of predicted lineage-specific regulatory elements across concurrent sequence-to-function studies.**

(**a**) Venn diagram illustrating the intersection of genomic loci predicted to have human-specific regulatory activity across three studies: Hominin Gain-linCREs (this study, dark blue), human-specific cerebellar *cis*-regulatory elements (Sarropoulos et al., dark orange), and human predicted increased chromatin accessibility regions (hPICAs, Wang et al., light orange).

(**b**) Heatmap displaying the significance of overlap enrichments ($-\log_{10}$(pAdj)) between Hominin Gain-linCREs stratified across 100 Enformer biosample contexts (rows) and hPICAs stratified across 111 cell types (columns).
